# Mesenchymal stem cells suppress leukemia via macrophage-mediated functional restoration of bone marrow microenvironment

**DOI:** 10.1101/661298

**Authors:** Chengxiang Xia, Tongjie Wang, Hui Cheng, Yong Dong, Qitong Weng, Guohuan Sun, Peiqing Zhou, Kaitao Wang, Xiaofei Liu, Yang Geng, Shihui Ma, Sha Hao, Ling Xu, Yuxian Guan, Juan Du, Xin Du, Yangqiu Li, Xiaofan Zhu, Yufang Shi, Sheng Xu, Demin Wang, Tao Cheng, Jinyong Wang

**Author notes:** Correspondences: Jinyong Wang (J.W.), Tao Cheng (T.C.). Equal contributors (Chengxiang Xia, Tongjie Wang, Hui Cheng).

## Abstract

Bone marrow (BM) mesenchymal stem cells (MSCs) are critical components of the BM microenvironment and play an essential role in supporting hematopoiesis. Dysfunction of MSCs is associated with the impaired BM microenvironment that promotes leukemia development. However, whether and how restoration of the impaired BM microenvironment can inhibit leukemia development remain unknown. Using an established leukemia model and the RNA-seq analysis, we discovered functional degeneration of MSCs during leukemia progression. Importantly, intra-BM instead of systemic transfusion of donor healthy MSCs restored the BM microenvironment, thus systemically altering cytokine expression patterns, improving normal hematopoiesis, reducing tumor burden, and ultimately prolonging survival of the leukemia-bearing mice. Donor MSC treatment restored the function of host MSCs and reprogrammed host macrophages to fulfill tissue-repair function. Transfusion of MSC-reprogrammed macrophages largely recapitulated the therapeutic effects of MSCs. Further, we found that donor MSCs reprogrammed macrophages to reduce leukemia burden through autocrine of IL-6. Taken together, our study reveals that donor MSCs reprogram host macrophages to restore the BM microenvironment and inhibit leukemia development, thus offering rationales for local MSC administration as a potentially effective therapy for leukemia.

**Key Points:** *Key Point 1*: Intra-BM transfusion of MSCs restores the BM microenvironment, improves thrombopoiesis, and suppresses MDS/MPN initiated by *Nras* mutation.

*Key Point 2*: Donor MSCs reprogram macrophages to restore the BM microenvironment, improve thrombopoiesis, and suppress leukemia.

## Introduction

BM MSCs are key components of the BM stromal microenvironment, regulating homeostasis of the stromal niche and hematopoiesis^1^. Accumulating evidence uncovers that dysfunction of MSCs is associated with leukemia progression. Leukemic MSCs in a mouse T-ALL model suppressed normal hematopoiesis^2^. Deletion of *Dicer1* gene in MSCs leads to impaired osteogenic differentiation, thrombopenia, and secondary tumorigenesis^3^. In myeloproliferative neoplasms, the leukemic cells reprogram MSCs to support proliferation of leukemic stem cells^4^. Ablation of MSC-derived osteoblasts impairs the homeostasis of hematopoietic stem cells (HSC) and accelerates leukemogenesis in chronic myeloid leukemia^5^. In a myelodysplastic/myeloproliferative neoplasms (MDS/MPN) model, *Ptpn11* mutation in MSCs hyperactivates HSC by secreting CCL3, systemically resulting in myeloid-biased proliferation and promoting leukemia progression^6^. Chromosomal abnormalities in MSCs occur in over 16% MDS/acute myeloid leukemia (AML) patients, which leads to a shorter survival^7^. MSCs of every MDS patient show features of stemness loss, aging, and osteogenic differentiation defect, which results in insufficient hematopoiesis^8^. In a patient-derived xenograft (PDX) model, only co-transplantation of tumor cells and the same patient-derived MSCs successfully recapitulates the MDS phenotype, indicating that besides the loss of function of supporting hematopoiesis, MSCs in MDS patients may acquire a function of preferentially promoting tumorigenesis^9^. Despite these findings, the causal relationship between tumor cells and the BM microenvironment during leukemia development and progression remains elusive.

MSC therapy has been widely used in treating immune-related graft-vs-host disease (GVHD) and inflammation-related diseases^10, 11^. Over the decades, a lot of evidence demonstrates that MSCs regulate innate and adaptive immune responses largely by secreting distinct sets of cytokines, growth factors, and chemokines depending on different disease contexts^12–16^. Given the short lifespan of donor MSCs after transfusion^17^, the underlying molecular and cellular mechanisms by which these cells produce therapeutic effects remain elusive. It is also completely unknown whether donor MSCs can restore the impaired BM microenvironment and consequently suppress disease progression in leukemia setting.

Macrophages are pivotal for maintenance of the tissue microenvironment, tissue repair, and even the tumor microenvironment^18–22^. BM resident macrophages maintain the homeostasis of HSCs and loss of these macrophages leads to mobilization of HSCs into peripheral blood (PB)^23^. The functions of macrophages are plastic and can be reshaped by distinct sets of soluble factors. When performing tissue repair, macrophages highly express arginase 1 (*Arg1*)^24^, an enzyme that converts L-arginine to urea and L-ornithine. After co-culture with MSCs, macrophages can be polarized from pro-inflammation (M1) to anti-inflammation (M2) type, up-regulating IL-10 and CD206 and down-regulating IL-6 and IL-1β^25^. Upon stimulated by LPS or TNF-α, MSCs can cross-talk with lung macrophages and reprogram these macrophages to secrete IL-10 to alleviate sepsis^26^. Despite these knowledge, whether healthy MSCs can reprogram macrophages from leukemia-bearing host to repair the damaged BM microenvironment is not known.

Using the established mouse model mimicking chronic MPN/MDS diseases^27–29^, we discovered that the deteriorating BM microenvironment was associated with disease progression. Intra-BM instead of systemic transfusion of healthy MSCs restored the local BM microenvironment, improved thrombopoiesis, reduced tumor burden, and prolonged survival of leukemia-bearing mice. Mechanistically, we found that MSCs suppress leukemia development through resident macrophages and autocrine effect of IL-6. Our study demonstrates that intra-BM transfusion of MSCs can restore the local BM niche to systemically prevent leukemia progression and can be a novel therapy for leukemia.

## Materials and Methods

### Mice

All mouse strains were maintained on C57BL/6 genetic background. Mice expressing the conditional oncogenic NrasG12D mutation (a gift from Dr. Jing Zhang lab at University of Wisconsin-Madison, Wisconsin, USA) were crossed to Vav-Cre mice to generate *LSL Nras*/*+*; *Vav-Cre* compound mice (NV mice). Genotyping of the adult mice was performed as described previously^27^. Vav-Cre strain (CD45.2), wild-type CD45.2, CD45.1 strain (C57BL/6) and *Il6* knock out strain (*Il6*^-/-^, C57BL/6) were purchased from Jackson lab. GFP strain (CD45.2) was gifted by Guangdong Laboratory Animals Monitoring Institute. *MLL-AF9* AML model mice were maintained a specific pathogen-free animal facility at the State Key Laboratory of Experimental Hematology. All mice were maintained within the SPF grade animal facility of Guangzhou Institution of Biomedicine and Health, Chinese Academy of Science (GIBH, CAS, China). All animal experiments were approved by the Institutional Animal Care and Use Committee of Guangzhou Institutes of Biomedicine and Health (IACUC-GIBH).

### NrasG12D leukemia model

White blood cells (CD45.2^+^, 0.3 million) after depletion of stromal cells from NrasG12D compound mice (*LSL Nras/+; Vav-Cre*) or control mice (CD45.2 strain) were sorted and transplanted into sublethally (6.5 Gy, RS2000, Rad Source Inc) irradiated CD45.1 recipient by retro-orbital intravenous injection. Mice were fed with trimethoprim-sulfamethoxazole-treated water for two weeks to prevent infection. Hematopoietic lineages in PB were assessed monthly by flow cytometry. During the development of NrasG12D-induced leukemia, the CD11b^+^ percentage in PB indicated the tumor burden (CD11b^+^%).

### RNA-Seq and data analysis

For MSC library preparation, MSCs were sorted from wild type or leukemia-bearing mice, and recovered MSCs were sorted from leukemia-bearing mice 8 weeks post treatment with GFP^+^ donor MSCs. MSCs were sorted from two mice of each group. 1000 target cells per sample were sorted into 500 µl DPBS-BSA buffer (0.5%BSA) using 1.5ml EP tube and transferred into 250 µl tube to spin down with 500 g. The cDNA of sorted 1000-cell aliquots were generated and amplified as described previously^30^. The qualities of the amplified cDNA were examined by Q-PCR analysis of housekeeping genes (*B2m, Actb, Gapdh, Ecf1a1*). Samples passed quality control were used for sequencing library preparation by illumina Nextera XT DNA Sample Preparation Kit (FC-131-1096).

For macrophages (*in vivo*) library preparation, macrophages were sorted from BM of leukemia-bearing mice before or after MSC treatment (12 hours post MSC treatment). Macrophages were also sorted after 12 hours of co-culture with MSCs. 1 × 10^5 target cells per sample were sorted and total RNA was extracted using the RNeasy micro kit with on-column DNase treatment (Qiagen, 74004) according to manufacture’s protocol. cDNA library was constructed using VAHTSTM mRNA-seq V3 Library Prep Kit for Illumina (Vazyme, NR611) according to manufacture’s protocol. The qualities of the cDNA were examined by qPCR analysis of housekeeping genes (*B2m, Actb, Gapdh, Ecf1a1*). Samples that passed quality control were used for sequencing.

For data analysis, all libraries were sequenced by illumina sequencers NextSeq 500. The fastq files of sequencing raw data samples were generated using illumina bcl2fastq software (version: 2.16.0.10) and were uploaded to Gene Expression Omnibus public database (GSE 125029). Raw reads were aligned to mouse genome (mm10) by HISAT2^31^ (version: 2.1.0) as reported. And raw counts were calculated by featureCounts of subread^32^ (version 1.6.0). Differential gene expression analysis was performed by DESeq2^33^ (R package version: 1.18.1). Unsupervised clustering analysis was performed using facotextra (R package, version: 1.0.5). Heatmaps were plotted using gplots (R package, version 3.01). GSEA was performed as described^34^, and gene-ontology (GO)-enrichment analysis were performed by clusterProfiler^35^ (R package, version: 3.6.0). MSC stemness related genes and MSC osteogenesis related genes for heatmaps were from references as follows: MSC stemness-related genes^36–38^ and MSC osteogenesis-related genes^36, 39^. The gene sets for GSEA were from literatures as follows: angiogenesis-related genes in macrophages^40^, cell migration-related genes in macrophages (from MSigDB genesets), and secreted factors by macrophages^41, 42^.

### MSC treatment for leukemia-bearing mice

For MSC transfusion, multiple approaches including retro-orbital, tail intravenous, and local intra-BM transfusion were applied independently. For tail vein transfusion, each leukemia-bearing mouse was injected with 2.5 × 10^7 MSCs/kg (Passage 2) in 100 μl DPBS by tail vein transfusion. For retro-orbital transfusion, each leukemia-bearing mouse was injected with 2.5 × 10^7 MSCs/kg (Passage 2) in 200 μl DPBS by retro-orbital transfusion. For local intra-BM transfusion, tibia of each leukemia-bearing mouse was injected with 2.5 × 10^7 MSCs/kg (Passage 2) in 20 μl DPBS by local intra-BM transfusion. MSCs were injected once every two weeks and continued in a time window of 16 weeks. Every tibia was treated once per month by switching the injection site every other dose. The control mice were injected with DPBS following the same treatment procedure as MSCs. Analysis of platelets and CD11b^+^ cells in PB was performed monthly.

### GFP-MSCs and BM macrophage co-culture assay

Short-term co-culture assay was performed, with each well containing: 1 × 10^5 GFP-MSCs (passage 2; healthy MSCs were isolated from GFP mice, *Il6*^-/-^ MSCs were isolated from *Il6*^-/-^ mice) and 2 × 10^6 CD11b^+^ leukemic cells sorted from leukemia-bearing mice in 2 mL culture medium of α-MEM, 10% FBS and 50 ng/ml SCF. MSCs and CD11b^+^ leukemic cells were incubated either by direct-contact culture or transwell culture for 12 hours at 37°C under 5% CO_2_ in a humidified incubator. MSC-reprogrammed macrophages from leukemia-bearing mice (CD11b^+^F4/80^+^) were sorted for detecting the gene expression by Q-PCR.

### Treatment for leukemia-bearing mice with MSC-reprogrammed macrophages

1 × 10^5 MSCs were seeded into each well of six-well plates. CD11b^+^ leukemic cells were enriched from BM of leukemia-bearing mice with severe tumor burden (CD11b^+^% in PB > 60%). Then 2 × 10^6 CD11b^+^ leukemic cells were directly co-cultured with MSCs. After 12 hours, macrophages were sorted for transfusion. Leukemia-bearing mice with severe tumor burden were treated by intra-BM transfusion of PBS or MSC-reprogrammed macrophages from leukemia-bearing mice (E-Mac). A dose of 1 million macrophages/mouse in 20 μl PBS were delivered into the tibia cavity using 29-gauge needle. Every tibia was treated once per two weeks by switching the injection site every other dose. Analysis of platelets and CD11b^+^ cells in PB was performed monthly.

### Statistical analysis

The data were represented as mean ± SD. Two-tailed independent Student’s t-tests were performed for comparison of two groups of data (SPSS v.23, IBM Corp., Armonk, NY, USA). For the analysis of three groups or more, one-way ANOVA was used (SPSS v.23, IBM Corp., Armonk, NY, USA), and further significance analysis among groups was analyzed by Post Hoc Test (equal variances, Turkey-HSD; unequal variances, Games-Howell). Kaplan-Meier method was used to calculate survival curves of leukemia, and Log-rank (Mantel-Cox) test was performed to compare differential significance in survival rates. P values of less than 0.05 were considered statistically significant (*p < 0.05, **p < 0.01, ***p < 0.001).

## Results

### Deterioration of BM MSCs accompanies the development of *Nras*-mutant-induced leukemia

Mice carrying an endogenous mutant *Nras* allele develop myelodysplastic/myeloproliferative neoplasms (MDS/MPN)-like leukemia with a long latency^27–29, 43^. Here we found the primary BM leukemic cells failed to accelerate the disease in the secondary recipient mice, implying a role of the BM microenvironment in disease etiology (Figure S1). We hypothesized that the BM microenvironment is impaired by *Nras*-mutant leukemic cells, which in return impedes normal hematopoiesis and accelerates leukemia progression. Indeed, we observed quantitative decreases and functional degeneration of MSCs (Ter119^-^CD45^-^CD31^-^Sca1^+^CD51^+^CD146^+^) during disease development and progression (Figure 1A-C, and Figure S2). To further characterize the residual MSCs in mice with leukemia, we performed RNA-Seq analysis of the residual MSCs from leukemia-bearing mice at an early disease phase (CD11b^+^% in PB: 35%-45%). Consistent with the quantitative and functional reduction, the expression of the transcription factor *Gnl3*^36^, an indicator of MSC self-renewal, was significantly down-regulated in MSCs from leukemia-bearing mice relative to wild-type mice (Figure 1D). The expression of *Nt5e* (CD73)*, Thy1* (CD90), *Vcam1* (CD106)*, Cd81*, *Sdc4, Itgb1* and *Anpep*^36–38^, encoding surface markers on three-lineage-potent MSCs but not on uni-lineage-primed MSCs, was markedly reduced in MSCs from leukemia-bearing mice (padj < 0.05, fold change > 1.6) (Figure 1D). Furthermore, the expression of *Bgn*, *Bmp4*, *Col1a1*, *Csf1*, *Dcn*, *Dkk2*, *Mmp13*, *Ogn*, *Wisp1*, and *Wisp2*^36, 39^, pivotal for osteogenic differentiation, was markedly suppressed in the residual MSCs (padj < 0.05, fold change > 2) (Figure 1E). *In vitro* differentiation assay confirmed that osteogenic and adipogenic differentiation potential in leukemic MSCs were impeded (Figure S3). MSCs fulfill their tissue-specific and condition-responsive regulatory functions through secreting distinct types of soluble factors^44^. Under leukemia condition, the residual MSCs indeed secreted much less soluble factors, including *Il6, Il11, Ccl2, Ccl7, Cxcl12, Cxcl13* and *Cxcl14* (padj < 0.05, fold change > 2), compared to MSCs from normal wild-type mice (Figure 1F). These molecules are pivotal for tissue repairing^45–50^. Indirect immunofluorescence assay and intracellular flow cytometry staining confirmed that the levels of IL-6 (Figure 1G-H), CCL2 (Figure 1I-J), and CXCL12 (Figure 1K-L) proteins in leukemic MSCs were significantly lower than healthy MSCs (p < 0.001). In addition, *Ccl5,* a chemokine involved in the pathogenesis of MPN^51^ and the inhibition of thrombopoiesis^52^, was significantly up-regulated in the residual MSCs from leukemia-bearing mice (padj < 0.05, fold change > 2). Gene set enrichment analysis (GSEA) further revealed features of inflammation in the residual MSCs (Figure S4). Collectively, these results show that the MSCs dramatically deteriorate during the disease development and progression of *Nras*-mutation-caused leukemia.

**Figure 1:**
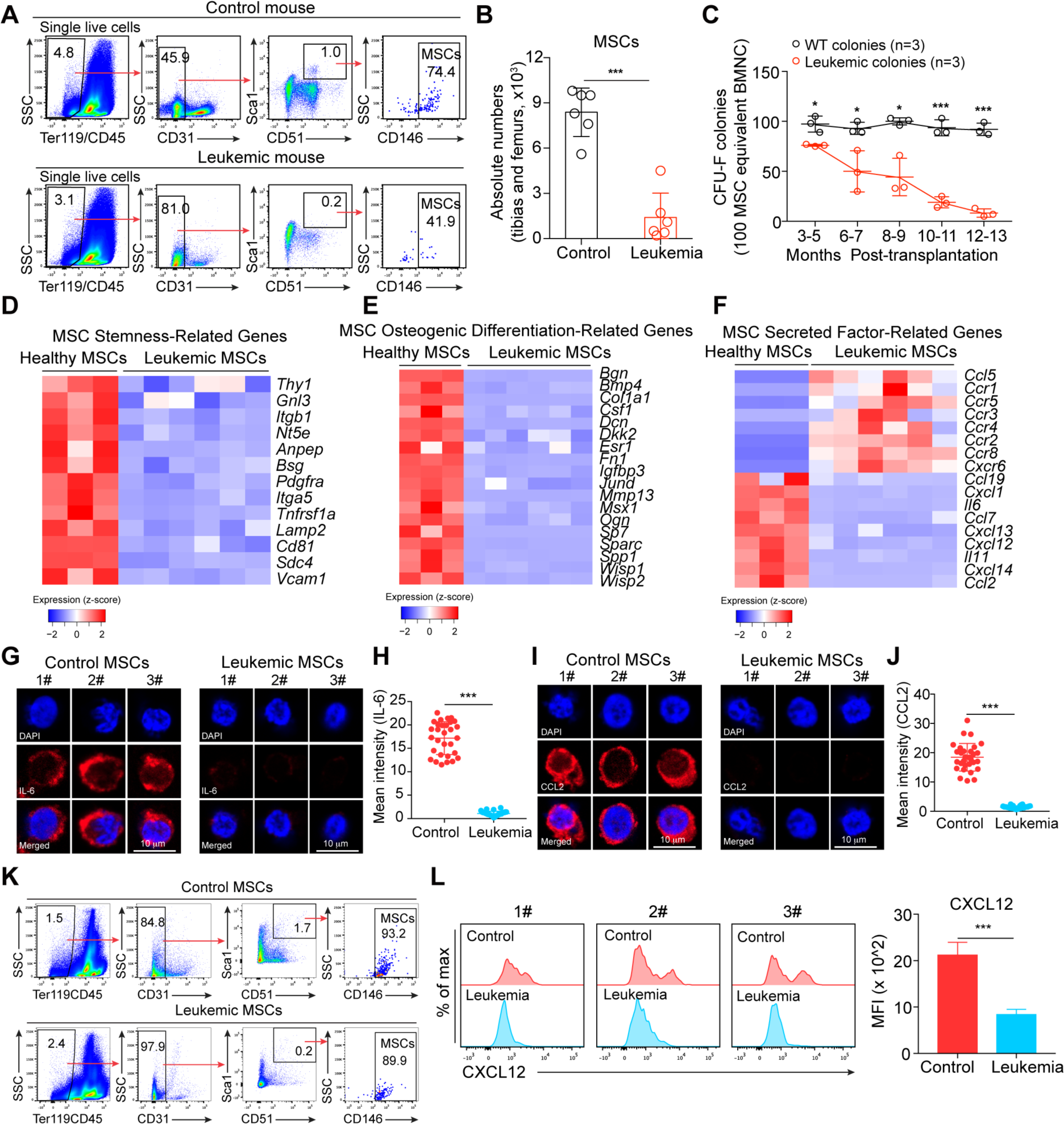
Impaired BM MSCs in mice with NrasG12D mutation-induced leukemia. CD45.2^+^ BMNC from LSL *Nras*/+; *Vav-Cre* mice (NV mice) were sorted and transplanted into sublethally irradiated (6.5 Gy) individual recipients (CD45.1 strain) with a cell dose of 0.3 million per recipient. For control recipient mice (CD45.1 strain), 0.3 million sorted CD45.2^+^ BMNC from WT mice were transplanted. **(A)** Gating strategies for BM MSCs. MSCs are defined as Ter119^-^ CD45^-^CD31^-^Sca1^+^CD51^+^ CD146^+^. Plots from one representative control mouse (CD11b^+^% in PB = 10%) and one leukemia-bearing mouse (CD11b^+^% in PB > 60%) of six mice of each group are shown. The nucleated cell mixtures of BM and compact bones were prepared for flow cytometry analysis of MSCs. **(B)** Statistical analysis of the absolute numbers of MSCs in tibias and femurs from control and leukemia-bearing mice. Data are analyzed by unpaired student’s t-test (two-tailed). ***p < 0.001. Data are represented as mean ± SD (n = 6 mice for each group from multiple independent experiments). **(C)** Kinetic analysis of functional MSCs in CFU-F assay. Total BMNC equivalent to 100 MSCs calculated by the percentages of MSCs in individual BMNC samples by flow cytometry analysis. Data are analyzed by unpaired student’s t-test (two-tailed). *p < 0.05, ***p < 0.001. For each time point, three independent experiments were performed. Data are represented as mean ± SD. **(D)** Heatmaps of MSC stemness-related genes differentially expressed between healthy MSCs and MSCs from leukemia-bearing mice (padj < 0.05, fold change > 1.6). One thousand sorted MSCs (Ter119^-^CD45^-^CD31^-^Sca1^+^CD51^+^) from the nucleated cell mixtures of BM and compact bones of independent leukemia-bearing mice, control mice, and MSC-treated leukemia-bearing mice were used as each cell sample input for RNA-Seq. Leukemia-bearing mice (CD11b^+^% in PB = 35%-45%) and control mice (CD11b^+^% in PB = 10-15%) were accumulated from multiple independent experiments. The expression value (DESeq2 normalized counts) of each gene was converted to z-score values (red, high; blue, low), and the heatmaps were plotted by gplots (heatmap.2). Columns represent the indicated cell subsets in nine MSC samples (Healthy MSCs from control mice: n = 3, MSCs from leukemia-bearing mice: n = 6). **(E)** Heatmaps of osteogenic differentiation-related genes differentially expressed between healthy MSCs and MSCs from leukemia-bearing mice (padj < 0.05, fold change > 2). The expression value (DESeq2 normalized counts) of each gene was converted to z-score values (red, high; blue, low), and the heatmaps were plotted by gplots (heatmap.2). Columns represent the indicated cell subsets in nine MSC samples (Healthy MSCs: n = 3, MSCs from leukemia-bearing mice: n = 6). **(F)** Heatmaps of MSC secreted factor-related genes differentially expressed between healthy MSCs and MSCs from leukemia-bearing mice (padj < 0.05, fold change > 2). The expression value (DESeq2 normalized counts) of each gene was converted to z-score values (red, high; blue, low), and the heatmaps were plotted by gplots (heatmap.2). Columns represent the indicated cell subsets in nine MSC samples (Healthy MSCs: n = 3, MSCs from leukemia-bearing mice: n = 6). **(G)** Single-cell imaging of intracellular IL-6 proteins by indirect immunofluorescence assay (IFA) in primary MSCs sorted from the control and leukemic mice. Images of three single representative cells of each group are shown. **(H)** Statistical analysis of mean intensities of IL-6 fluorescence in control and leukemic MSC samples. Each dot represents a single cell. Data are analyzed by unpaired student’s t-test (two-tailed). ***p < 0.001. Data are represented as mean ± SD. Control, n=30; Leukemia, n=30. Single-cell imaging of intracellular CCL2 proteins by IFA in primary MSCs sorted from the control and leukemic mice. Images of three single representative cells of each group are shown. Statistical analysis of mean intensities of CCL2 fluorescence in control and leukemic MSC samples. Each dot represents a single cell. Data are analyzed by unpaired student’s t-test (two-tailed). ***p < 0.001. Data are represented as mean ± SD. Control, n=30; Leukemia, n=30. **(K)** Representative intracellular staining plots of CXCL12 proteins in control and leukemic MSCs analyzed by flow cytometry. **(L)** Statistical analysis of mean fluorescence intensities (MFI) of CXCL12 in control and leukemic MSCs. Data are analyzed by unpaired student’s t-test (two-tailed). ***p < 0.001. Plots from three representative control mice and leukemia-bearing mice are shown. Data are represented as mean ± SD (n = 6 mice for each group).

### Intra-BM transfusion of healthy MSCs improves thrombopoiesis, reduces tumor burden and improves survival of the leukemia-bearing mice

We hypothesized that restoration of the impaired BM microenvironment in leukemia-bearing mice might suppress/delay the disease progression. To test this hypothesis, we attempted healthy MSC treatment using GFP-tagged MSCs isolated from the tibias and femurs of healthy mice as previously reported^53^. The isolated primary MSCs were expanded shortly *in vitro* to passage two (P2) and cryopreserved. For MSC treatment, the cryopreserved P2 MSCs were recovered and cultured for five days, phenotypically identified (CD45^-^Ter119^-^CD31^-^ CD51^+^CD105^+^LepR^+^PDGFRα^+^PDGFRβ^+^Sca1^+^) (Figure S5), and suspended in DPBS (2.5 × 10^7/ml) for transfusion. Initially, we adopted a direct delivery procedure by injecting donor MSCs every two weeks either via tail vein (dose: 0.5 × 10^6/mouse) (Figure S6A) or retro-orbital (dose: 0.5 × 10^6/mouse) transfusion (Figure S6B) into the leukemia-bearing mice at a late disease phase (CD11b^+^ cells > 60% in PB). However, these delivery approaches failed to produce therapeutic effects. *In vitro* cultured MSCs lose their natural homing feature^54^, the retro-orbital and intravenous injection of cultured MSCs failed to home to bone marrow (Figure S6C-D). Thus, we attempted intra-BM transfusion to overcome the homing defect caused by *in vitro* culture. A sequential doses of MSCs (2.5 × 10^7 MSCs/kg per dose in 20 µL DPBS) were injected into the tibia cavities of leukemia-bearing mice with two-week intervals for up to 16 weeks (Figure 2A). Strikingly, the tumor burden continuously decreased during MSC treatment (Figure 2B and Supplementary Figure S7). Consequently, the survival of treated leukemia-bearing mice was significantly prolonged (Untreated: 51.5 days, MSC-treated: >115 days, p < 0.001) (Figure 2C). Therefore, intra-BM transfusion of healthy donor MSCs improves the survival of leukemia-bearing mice.

**Figure 2:**
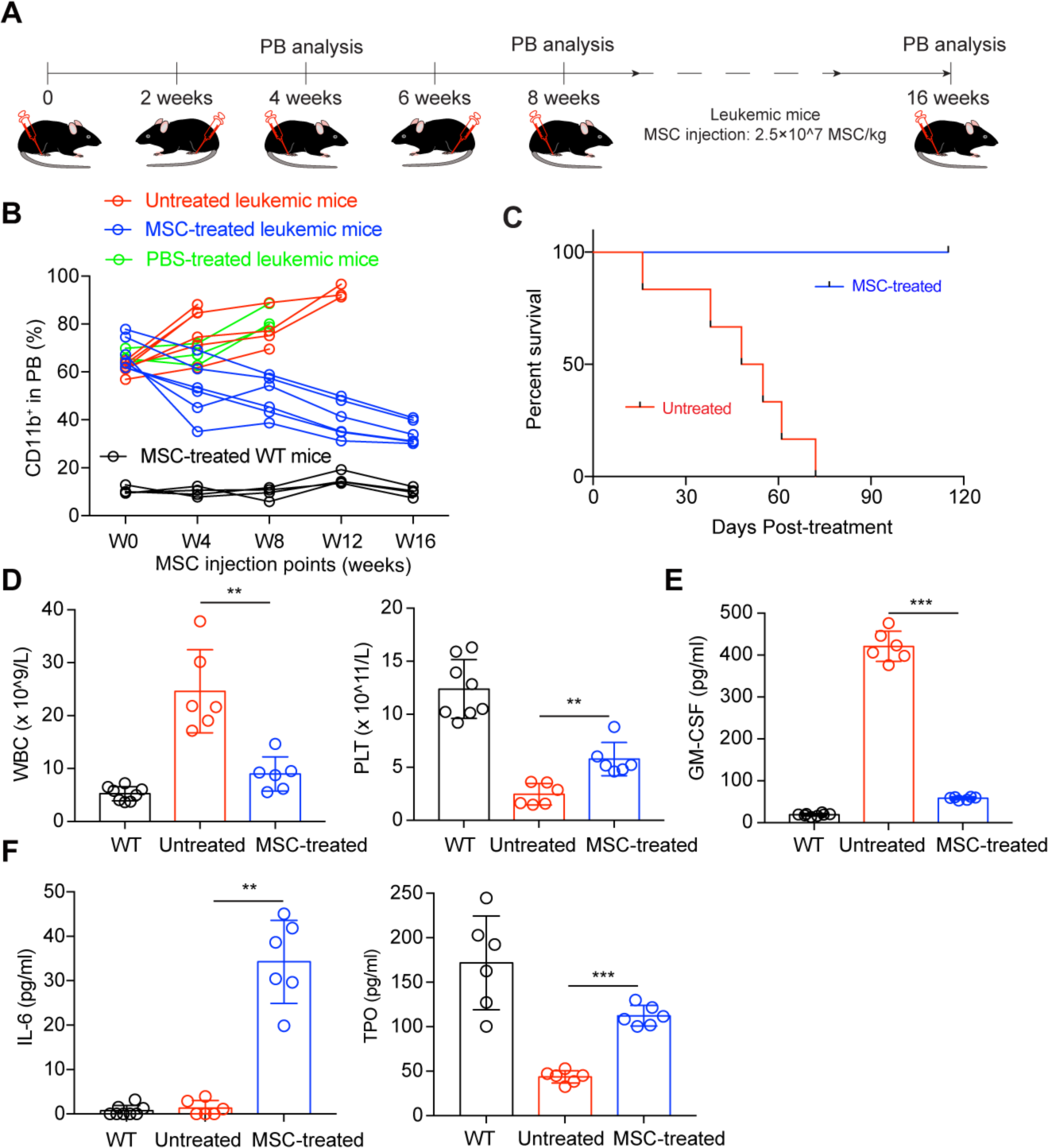
Intra-BM transfusion of donor MSCs prolongs survival of leukemia-bearing mice. **(A)** Schematic diagram of MSC transfusion strategy. The donor MSCs prepared for transfusion were isolated from the compact bone and BMNC of three to four-week-old healthy GFP mice. The isolated MSCs were expanded *in vitro*, and the secondary passage products were used for transfusion. Leukemia-bearing mice with severe tumor burden (CD11b^+^% in PB > 60%) were treated by intra-bone-marrow transfusion of donor MSCs. WT mice were treated by the same procedure as treatment control, and leukemia-bearing mice without MSC treatment were used as untreated controls. A sequential doses of MSCs (2.5 × 10^7 MSCs/kg per dose in 20 µL DPBS) were delivered into the tibia cavity using 29-gauge needle. Every tibia was treated once per month by switching the injection site every other dose. **(B)** Kinetic analysis of tumor burden (CD11b^+^) of MSC-treated leukemia-bearing mice. The time window of MSC treatment is from 0 weeks (W0, treatment starting time) to 16 weeks (W16). Flow cytometry analysis of tumor burden (CD11b^+^) in PB was performed monthly. These leukemia-bearing mice were accumulated from multiple independent experiments. The individual leukemia-bearing mice were selected to perform treatment once they reached the leukemic burden standard (CD11b^+^% > 60% in peripheral blood), but at different time from multiple experiments. Untreated leukemia-bearing mice were used as disease control (red line), PBS-treated leukemia-bearing mice were used as injected control (green line), and MSC-treated WT mice were used as treatment control (black line). Untreated leukemia-bearing mice: n = 6; PBS-treated leukemia-bearing mice: n = 3; MSC-treated leukemia-bearing mice (blue line): n = 6; MSC-treated WT mice: n = 4. **(C)** Kaplan-Meier survival of MSC-treated leukemia-bearing mice. Kaplan-Meier survival curves of untreated (n = 6, Median survival = 51.5 days) and MSC-treated (n = 6, Median survival = 115 days) leukemia-bearing mice are shown. MSC treatment was terminated after 16 weeks. The untreated leukemia-bearing mice from the same batch were used as control (red line). Log-rank (Mantel-Cox) test: p < 0.001. **(D)** Statistical analysis of white blood cells (WBC) and platelets (PLT) counts in PB of WT mice, untreated leukemia-bearing mice, and MSC-treated leukemia-bearing mice at week nine since the MSC treatment. Data are analyzed by one-way ANOVA test. **p < 0.01. Data are represented as mean ± SD (n =6-8 mice for each group). **(E)** ELISA of GM-CSF levels in PB serum. Serum prepared from 200 μL PB of individual WT mice (n=8), untreated leukemia-bearing mice (Untreated, n=6) and MSC-treated leukemia-bearing mice (MSC-treated, n=6) eight weeks post MSC treatment. Data are analyzed by one-way ANOVA test. ***p < 0.001. Data are represented as mean ± SD. **(F)** ELISA of IL-6 and TPO levels in PB serum. Serum prepared from 200 μL PB of individual WT mice (n=6-8), untreated leukemia-bearing mice (Untreated, n=6) and MSC-treated leukemia-bearing mice (MSC-treated, n=6) eight weeks post MSC treatment. Data are analyzed by one-way ANOVA. **p < 0.01, ***p < 0.001. Data are represented as mean ± SD.

### MSC-treatment systemically re-balances myelopoiesis and activates megakaryopoiesis

We next investigated the underlying mechanisms associated with the systemically decreased tumor burden. We found that the hematopoiesis in the MSC-treated leukemia-bearing mice was re-balanced, demonstrated by significant decreases of white blood cells (Untreated vs. MSC-treated: 23.04 vs 8.876, p = 0.009), and significant elevation of platelets (Untreated vs. MSC-treated: 2.64 vs. 6.01, p = 0.004) (Figure 2D) in PB. On the contrary, the PBS-treated leukemia-bearing mice exhibited neither improved hematopoiesis (Figure S8) nor prolonged survival. High GM-CSF levels in serum are associated with the tumor burdens of CMML in patients^55^ and mouse models^27^. We analyzed the GM-CSF levels in serum of leukemia-bearing mice with or without MSC-treatment. As expected, the GM-CSF levels in MSC-treated leukemia-bearing mice were significantly decreased (> 7 folds) (p < 0.001) (Figure 2E), which was associated with the reduction of GM-CSF secreting tumor cells (Figure S9). IL-6 promotes thrombopoiesis by increasing systemic TPO levels^56^. Consistently, the MSC-treated leukemia-bearing mice exhibited elevated IL-6 (> 25 folds) and TPO (> 2 folds) levels (Figure 2F) in PB serum. Collectively, these results indicate that intra-BM transfusion of healthy donor MSCs systemically improves hematopoiesis and prolongs the survival of leukemia-bearing mice.

To further investigate the systemic effects of the local MSC-treatment on hematopoiesis in leukemia-bearing mice, we analyzed the ratios of myeloid progenitor subpopulations in MSC-treated leukemia-bearing mice. Consistent with the elevated platelet levels and reduced myeloid cells in PB, the MSC-treated leukemia-bearing mice showed increased proportions of megakaryocyte-erythroid progenitors (MEP) (> 1.6 folds) (p < 0.001) and decreased ratios of granulocyte-macrophage progenitors (GMP) (> 1.5 folds) (p < 0.001) in both injected and non-injected sites than those sites in PBS-treated leukemia-bearing mice (Figure 3A-B, Figure S10A). In addition, we observed increased (> 1.3 folds) ratios of mature megakaryocytes (≥ 8N) in both injected and non-injected sites in MSC-treated leukemia-bearing mice (Figure 3C-D, Figure S10B) in comparison with PBS-treated leukemia-bearing control mice (p < 0.001). Thus, these data demonstrate that MSC-treatment systemically re-balances myelopoiesis and activates megakaryopoiesis in leukemia-bearing mice.

**Figure 3:**
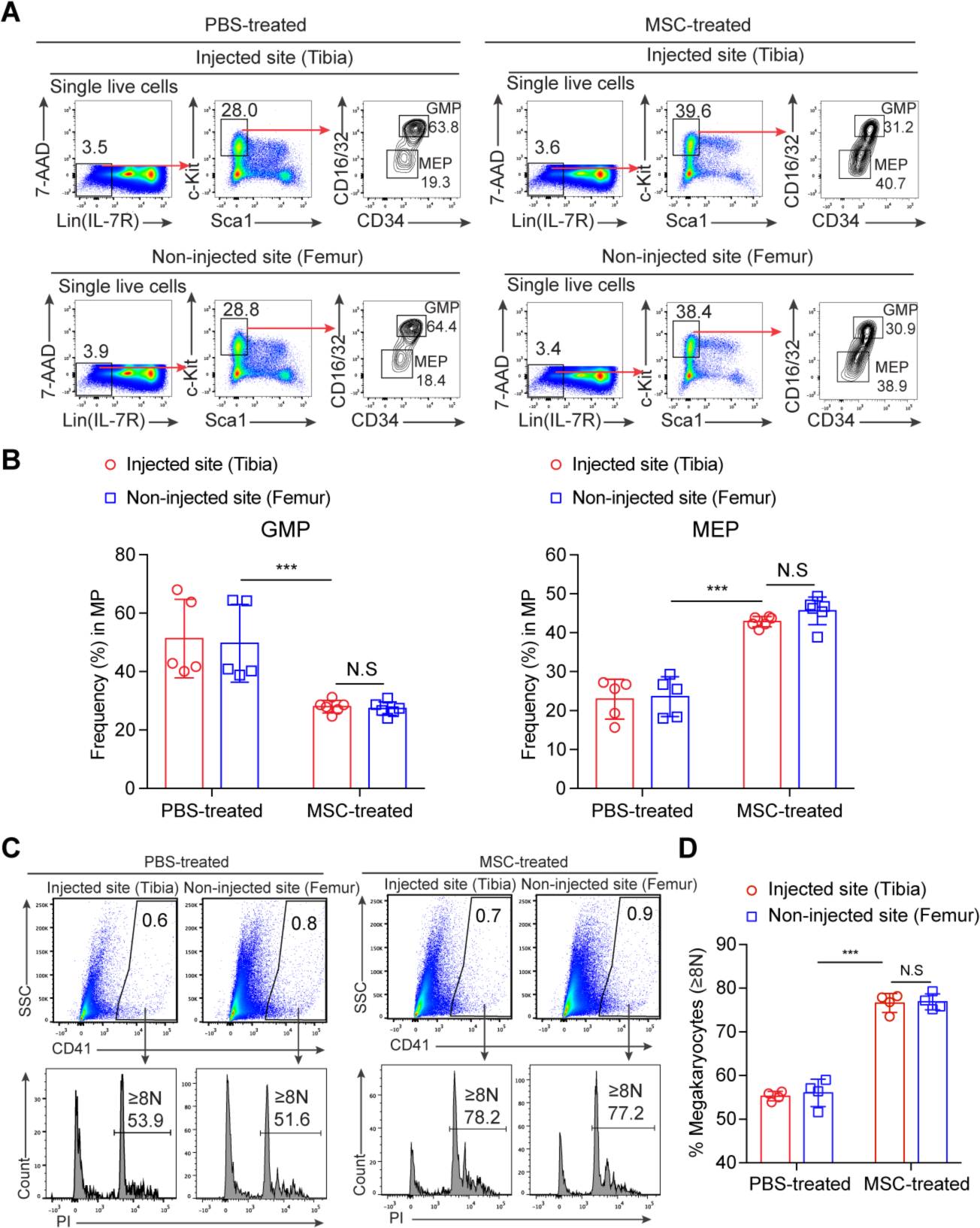
Systemically re-balanced myeloid lineage progenitor cells and systemically activated megakaryocytes in MSC-treated leukemia-bearing mice. **(A)** Ratios of myeloid progenitor subpopulations in MSC- and PBS-treated leukemia-bearing mice. Total bone marrow nucleated cells were from the MSC-injected site (tibia) and non-injected site (femur) of each leukemia-bearing mouse, or PBS-injected site (tibia) and non-injected site (femur) of each leukemia-bearing mouse four weeks post MSC/PBS treatment. GMP (granulocyte/macrophage progenitors): Lin^-^IL-7R^-^Sca1^-^c-Kit^+^CD34^+^CD16/32^high^; MEP (megakaryocyte/erythroid progenitors): Lin^-^IL-7R^-^Sca1^-^c-Kit^+^CD34^-^CD16/32^-^. **(B)** Statistical analysis of myeloid progenitor components (GMP and MEP) in the injected site (tibia) and non-injected site (femur) of each PBS-treated leukemia-bearing mouse and MSC-treated leukemia-bearing mouse four weeks post-MSC treatment. Data are analyzed by one-way ANOVA test. ***p < 0.001. Data are represented as mean ± SD (n = 5-6 mice for each group accumulated from multiple independent experiments). **(C)** Activation analysis of megakaryocytes in MSC- and PBS-treated leukemia-bearing mice. Plots from one representative PBS-treated leukemia-bearing mouse and MSC-treated leukemia-bearing mouse four weeks post MSC treatment are shown. Percentages of mature megakaryocytes with 8N and greater ploidy (≥ 8N) are shown. **(D)** Statistical analysis of the percentages of mature megakaryocytes (≥8N). Data are analyzed by one-way ANOVA test. ***p < 0.001, N.S indicates not-significant. Data are represented as mean ± SD (n = 4 mice for each group accumulated from multiple independent experiments).

### Recovered host MSCs are functional as healthy counterparts

Consistent with an early report^57^, our data showed that MSCs indeed support normal hematopoiesis. Interestingly, MSCs also promoted the growth of leukemic cells *in vitro*, which suggests that donor MSCs indirectly inhibit the tumor development *in vivo*. To investigate whether the improved hematopoiesis is associated with restoration of the BM microenvironment, we analyzed the MSC-treated tibias eight weeks after MSC treatment. Interestingly, host MSCs (GFP negative) were partially recovered (Figure 4A-B), but restricted to the locally treated tibias (Figure S11). Functionally, the recovered host MSCs formed markedly more CFU-F colonies than the residual MSCs from untreated leukemia-bearing mice (> 3.8 folds) (p < 0.001) (Figure 4C and Figure S12). To characterize the recovered MSCs at the transcriptome level, we sorted the recovered MSCs for RNA-Seq analysis. Unsupervised hierarchical clustering analysis showed that the recovered MSCs clustered closer to healthy MSCs (Figure 4D). Further, the expression of cytokines and chemokines, including *Il6*, *Ccl2, Ccl7, Ccl19, Cxcl12, Cxcl13,* and *Cxcl14*, was restored in the recovered MSCs compared to that in MSCs from untreated leukemia-bearing control mice (padj < 0.05, fold change > 2) (p < 0.05) (Figure 4E). Consistently, indirect immunofluorescence assay and intracellular flow cytometry staining confirmed that the levels of IL-6, CCL2, and CXCL12 proteins in recovered MSCs (from the injected sites) were significantly improved to comparable levels as in healthy MSCs (p < 0.001) (Figure 4F-I and S13). Therefore, donor MSC-treatment results in local functional restoration of host MSCs.

**Figure 4:**
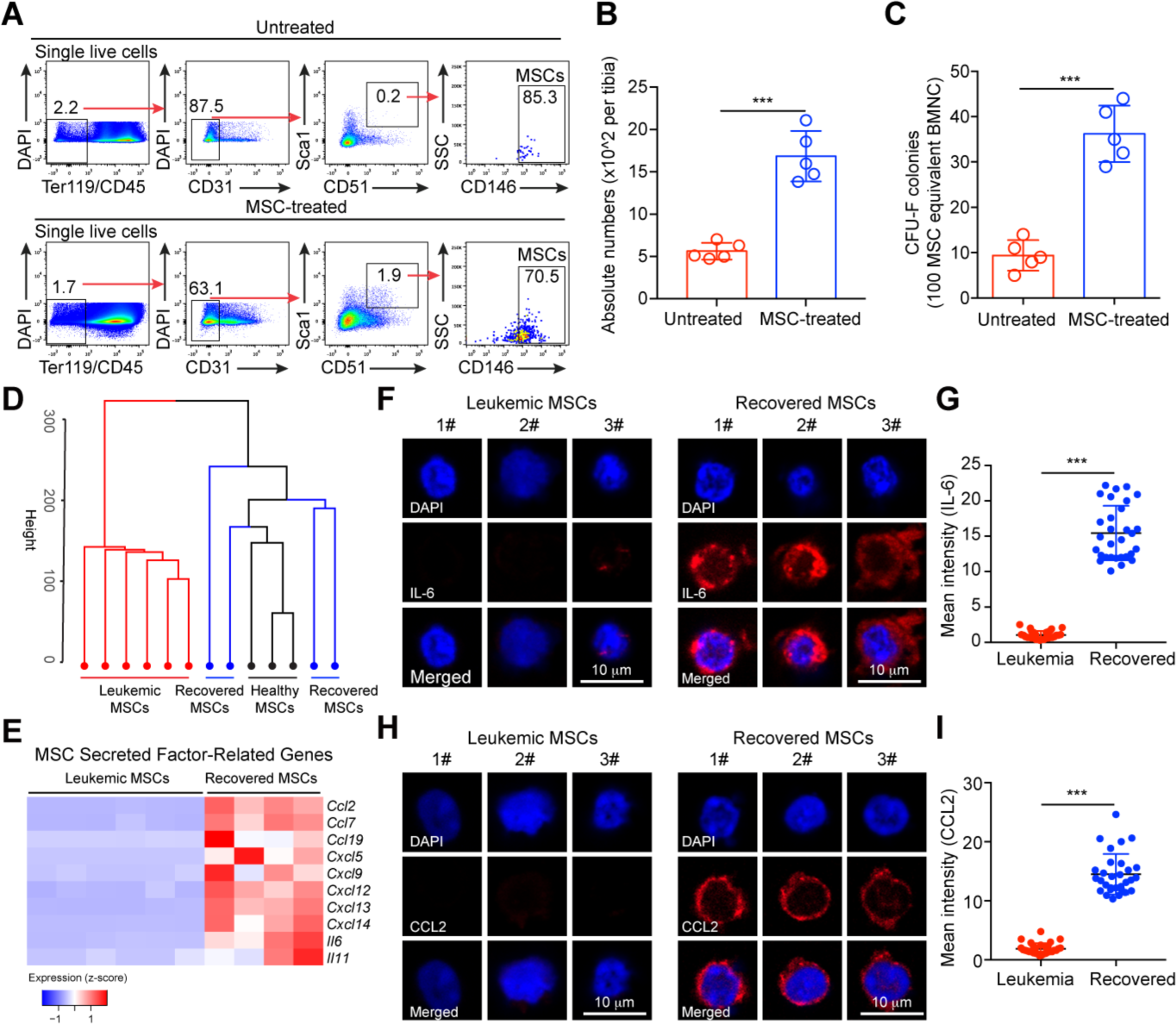
Characterization of recovered host MSCs from MSC-treated leukemia-bearing mice. **(A)** Flow cytometry analysis of recovered host MSCs in leukemia-bearing mice eight weeks post MSC treatment. Tibias of MSC-treated leukemia-bearing mice eight weeks after the first dose MSC treatment were analyzed. The leukemic mice are induced as described in Figure 1. The data of the MSC-treated tibias (injected sites) from MSC-treated leukemic mice and the control tibias from untreated leukemic mice are shown. Plots of one representative mouse from each group are shown. MSCs are defined as Ter119^-^CD45^-^CD31^-^Sca1^+^CD51^+^CD146^+^. The nucleated cell mixtures of BM and compact bones were prepared for flow cytometry analysis of MSCs. **(B)** Statistical analysis of the absolute numbers of host MSCs (GFP^-^, host-derived MSCs) in tibias from untreated and MSC-treated leukemia-bearing mice. Data are analyzed by unpaired student’s t-test (two-tailed). ***p < 0.001. Data are represented as mean ± SD (n = 5 mice for each group). **(C)** Statistical analysis of CFU-F colonies. The numbers of colonies of each group were counted after Giemsa staining. Data are analyzed by unpaired student’s t-test (two-tailed). ***p < 0.001. Data are represented as mean ± SD (n = 5 mice for each group). **(D)** Unsupervised hierarchical clustering of RNA-Seq data of MSCs from leukemia-bearing mice, healthy MSCs, and recovered MSCs (GFP^-^, recipient-derived). For each RNA-Seq sample, one thousand MSCs from leukemia-bearing mice, healthy mice, and MSC-treated leukemia-bearing mice were sorted and analyzed (n = 3-6). The raw reads (fastq files) from RNA-Seq were aligned to mouse genome by Tophat2 package, and further normalized by Cufflinks. Unsupervised hierarchical clustering was conducted by factoextra R package. **(E)** Heatmaps of MSC secreted factor-related genes differentially expressed between MSCs from leukemia-bearing mice and recovered MSCs (padj < 0.05, fold change > 1.4). The expression value (DESeq2 normalized counts) of each gene was converted to z-score values (red, high; blue, low), and the heatmaps were plotted by gplots (heatmap.2). Columns represent the indicated cell subsets in ten MSC samples (Leukemic MSCs: n = 6, Recovered MSCs: n = 4). **(F)** Single-cell imaging of intracellular IL-6 proteins by indirect immunofluorescence assay (IFA) in primary MSCs sorted from non-injected sites (Leukemic MSCs) and MSC-injected sites (Recovered MSCs) of MSC-treated leukemic mice. Images of three single representative cells of each group are shown. **(G)** Statistical analysis of mean intensities of IL-6 fluorescence in leukemic and recovered MSC samples. Each dot represents a single cell. Data are analyzed by unpaired student’s t-test (two-tailed). ***p < 0.001. Data are represented as mean ± SD. Control, n=30; Leukemia, n=30. **(H)** Single-cell imaging of intracellular CCL2 proteins by IFA in primary MSCs sorted from non-injected sites (Leukemic MSCs) and MSC-injected sites (Recovered MSCs) of MSC-treated leukemic mice. Images of three single representative cells of each group are shown. **(I)** Statistical analysis of mean intensities of CCL2 fluorescence in leukemic and recovered MSC samples. Each dot represents a single cell. Data are analyzed by unpaired student’s t-test (two-tailed). ***p < 0.001. Data are represented as mean ± SD. Control, n=30; Leukemia, n=30.

### The donor MSCs reprogram macrophages to execute tissue-repair function

We further investigated the cellular mechanism underlying the restored BM microenvironment mediated by donor MSCs under leukemia condition. BM macrophages play a pivotal role in maintaining the BM niche^18^. To study whether donor MSCs reprogram BM macrophages, we performed co-culture assay of healthy MSCs with BM macrophages (L-Mac) sorted from the leukemia-bearing mice *in vitro* for twelve hours and re-sorted the macrophages (E-Mac) for RNA-Seq analysis. GSEA illustrated that angiogenesis-related genes, including *Vegfa, Hif1a, Serpine1, Eng and Thbs1*^40^ (Figure S14A), were enriched among the differentially expressed genes in E-Mac (Figure 5A). Genes associated with cell migration, including *Sirpa* and *Ccl5*^58, 59^, were also enriched in E-Mac (Figure 5B and S14B). Further, gene-ontology analysis demonstrated features of positive regulation of cell migration and angiogenesis in E-Mac (Figure 5C). Elevated expression of genes encoding soluble factors, including *Tnf*, *Il1a*, *Ccr7*, *Ccl3*, and *Ccl5*, was also observed in E-Mac (Figure 5D). Indirect immunofluorescence assay confirmed that E-Mac had significantly higher levels of TNF-α, CCR7, and IL-1α proteins than L-Mac (p < 0.001) (Figure 5E and S15A). Furthermore, intracellular flow cytometry staining also showed that the levels of TNF-α and CCR7 proteins in L-Mac were significantly lower than E-Mac (p < 0.001) (Figure S15B-D). To characterize which subset of macrophages are involved in re-educating the bone marrow microenvironment, we analyzed the classical M1 and M2 macrophage subtypes both *in vitro* and *in vivo*. However, the results showed that MSC treatment caused no typical transition of M1 and M2 (Figure S16), which indicates a functional alteration of leukemic macrophages independent of M1 to M2 transition. RNA-Seq analysis showed that the expression of arginase 1 (*Arg1*), an indicator of tissue repair function^24^, was dramatically up-regulated over thousand folds in E-Mac (Figure 5F) after direct co-culture with MSCs *in vitro*. However, the *Arg1* expression in macrophages after transwell co-culture was barely elevated (Figure 5F), indicating that direct cell-cell interaction instead of MSC-secreted soluble factors is essential for the functional reprogramming. Consistent with the observation *in vitro*, the expression of *Arg1* was also significantly increased in BM macrophages directly isolated from MSC-treated leukemia-bearing mice (Figure 5G). Furthermore, intracellular flow cytometry staining confirmed that the ratios of Arg1^high^ macrophage subpopulation significantly increased in E-Mac at the MSC-treated sites (p < 0.01) (Figure 5H-I). Collectively, these results indicate that the donor MSCs reprogram BM macrophages from leukemia-bearing mice to execute tissue-repair function.

**Figure 5:**
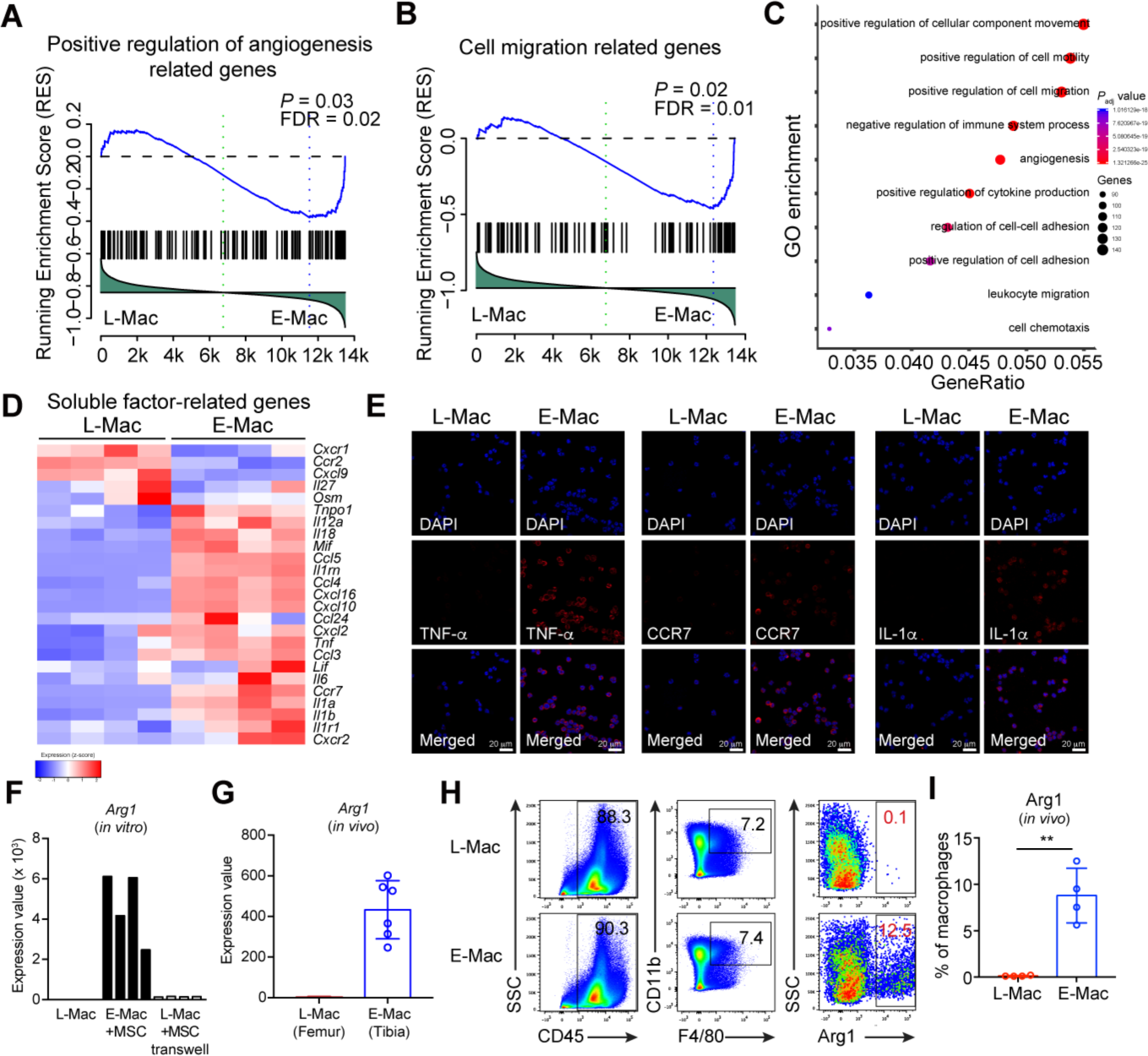
Characterization of MSC-reprogrammed BM resident macrophages isolated from leukemia-bearing mice. **(A)** Gene set enrichment analysis (GSEA) of the positive regulation of angiogenesis in L-Mac and E-Mac. L-Mac indicates leukemic macrophages. E-Mac indicates MSC-reprogrammed leukemic macrophages, which were co-cultured with MSCs *in vitro* for 12 h. DESeq2 normalized values of the expression data were used for GSEA analysis. **(B)** Gene set enrichment analysis (GSEA) of the cell migration-related genes in L-Mac and E-Mac. DESeq2 normalized values of the expression data were used for GSEA analysis. **(C)** Gene ontology (GO)–enrichment analysis of the 3277 differentially expressed genes between L-Mac and E-Mac: each symbol represents a GO term (noted in the plot); color indicates adjusted P value (padj (significance of the GO term)), and symbol size is proportional to the number of genes. **(D)** Heatmaps of soluble factor-related genes in MSC-reprogrammed leukemic macrophages. The expression value (DESeq2 normalized counts) of each gene was converted to z-score value (red, high; blue, low). The heatmaps were plotted by gplots (heatmap.2). Columns represent the indicated macrophage sample replicates (L-Mac: n = 4 biological replicates; E-Mac: n = 4 biological replicates). **(E)** Imaging of intracellular TNF-α, CCR7, and IL-1α proteins by indirect immunofluorescence assay (IFA) in sorted L-Mac (Non-injected site) and E-Mac (MSC-injected site) from MSC-treated leukemic mice 12 h post-treatment *in vivo*. The representative images of each group are shown. **(F)** RNA-Seq analysis of *Arg1* in leukemic macrophages co-cultured with MSCs *in vitro.* L-Mac indicates leukemic macrophages. E-Mac (+MSCs) indicates MSC-reprogrammed leukemic macrophages, which were co-cultured with MSCs *in vitro* for 12 h. L-Mac (+MSCs transwell) indicates leukemic macrophages, which were co-cultured in transwell with MSCs *in vitro* for 12 h. Y-axis indicates the expression value. The expression value (DESeq2 normalized counts) of each gene is illustrated by graphpad. Each column represents a replicate. **(G)** RNA-Seq analysis of *Arg1* in leukemic macrophages sorted from MSC-treated leukemic mice *in vivo*. Leukemic macrophages (CD11b^+^F4/80^+^) were sorted from the injected site (MSC-treated tibia) and non-injected site (untreated femur) of leukemic mice 12 h post-treatment. Y-axis indicates the expression value. The expression value (DESeq2 normalized counts) of each gene is illustrated by graphpad. Each column represents a replicate. **(H)** Representative intracellular staining plots of Arg1 proteins in MSC-reprogrammed leukemic macrophages (E-Mac, MSC-injected site) and leukemic macrophages (L-Mac, Non-injected site) from MSC-treated leukemic mice analyzed by flow cytometry. The ratios of Arg1^high^ macrophage subpopulation are shown from L-Mac and E-Mac. The bone marrow nucleated cells from leukemia-bearing mouse 12 h post-treatment were prepared for intracellular flow cytometry staining of Arg1. Macrophages are identified as CD45^+^CD11b^+^F4/80^+^. **(I)** Statistical analysis of the ratios of Arg1^high^ macrophage subpopulation in L-Mac and E-Mac. Data are analyzed by unpaired student’s t-test (two-tailed). **p < 0.01. Data are represented as mean ± SD (n = 4 mice for each group).

### The E-Mac treatment largely recapitulates the therapeutic effects of MSC treatment

Given the short lifespan of donor MSCs *in vivo*^17^, we speculated that MSCs mediate the restoration of the BM microenvironment of leukemia-bearing mice by reprogramming macrophages. We isolated macrophages from leukemia-bearing mice and co-cultured them with healthy MSCs for 12h, and then transplanted these E-Mac back into leukemia-bearing mice by intra-BM injection (Figure 6A). We indeed found that the thrombopoiesis was significantly improved (> 6 folds) after E-Mac treatment (p < 0.001) (Figure 6B-C). Host MSCs were also significantly increased (> 3 folds) in E-Mac-treated leukemia-bearing mice (p < 0.001) (Figure 6D-E). Consistent with the MSC treatment, we also observed increased ratios of mature megakaryocytes (p < 0.001) (Figure 6F-G) and alleviated tumor burden (CD11b^+^) (Figure S17) in E-Mac-treated leukemia-bearing mice. Collectively, these results demonstrate that MSC-reprogrammed macrophages largely recapitulate the therapeutic effects of MSCs.

**Figure 6:**
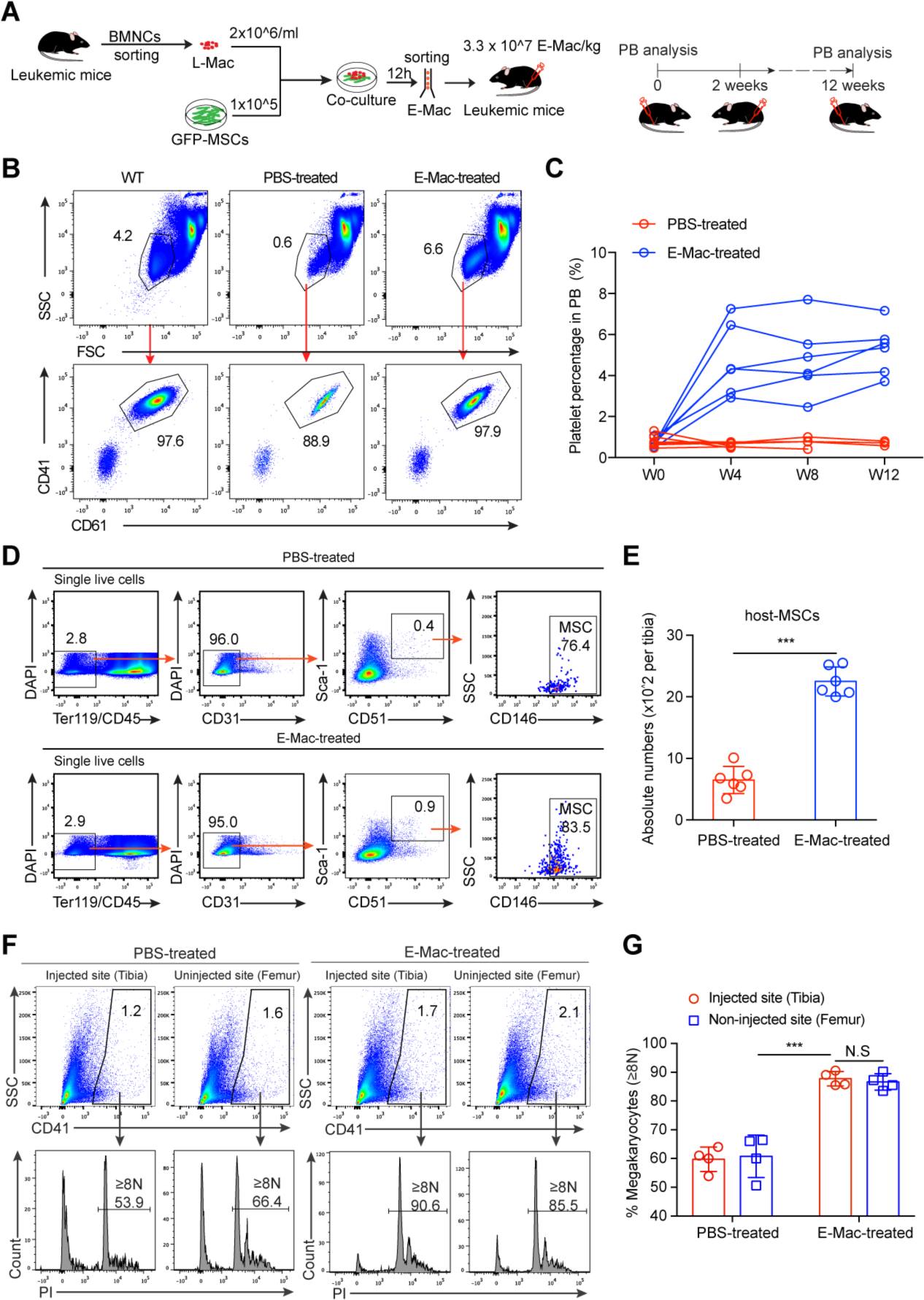
Intra-bone-marrow transfusion of MSC-reprogrammed macrophages largely rescues the therapeutic effects of MSC-treatment in leukemic mice. **(A)** Schematic diagram of MSC-reprogrammed macrophages transfusion strategy. 1 × 10^5 GFP^+^ MSCs were seeded into each well of six-well plates. CD11b^+^ leukemic cells were enriched from bone marrow of leukemic mice with severe tumor burden (CD11b^+^% in PB > 60%). Then 2 × 10^6 CD11b^+^ leukemic cells were directly co-cultured with MSCs. After 12 hours, leukemic macrophages (CD11b^+^F4/80^+^) were sorted for transfusion. Leukemic mice with severe tumor burden were treated by intra-bone-marrow transfusion of PBS or MSC-reprogrammed leukemic macrophages (E-Mac). A sequential doses of E-Mac (3.3 × 10^7 E-Mac/kg per dose in 20 μl PBS) were delivered into the tibia cavity using 29-gauge needle. Every tibia was treated once per two weeks by switching the injection site every other dose. Analysis of platelets and CD11b^+^ cells in PB was performed monthly. **(B)** Representative dot plots of platelet populations and quantitative gating, as identified by CD41 and CD61 staining in PB of WT mice and PBS/E-Mac treated leukemic mice. After 4 weeks of PBS/E-Mac treatment, PB of leukemic mice with PBS/E-Mac treatment was analyzed. **(C)** Kinetic analysis of platelets in PB of leukemia-bearing mice treated with PBS or E-Mac (n = 6 mice for each group). **(D)** Flow cytometry analysis of MSCs in leukemia-bearing mice post PBS/E-Mac treatment. Tibias of PBS/E-Mac treated leukemic mice at week five since the first dose of PBS/E-Mac treatment were analyzed. MSCs were defined as Ter119^-^CD45^-^CD31^-^Sca1^+^CD51^+^CD146^+^. The nucleated cell mixtures of BM and compact bones were prepared for flow cytometry analysis of MSCs. **(E)** Statistical analysis of the absolute numbers of host MSCs in tibias from PBS/E-Mac treated leukemic mice. Data are analyzed by unpaired student’s t-test (two-tailed). ***p < 0.001. Data are represented as mean ± SD (n = 6 mice for each group). **(F)** CD41^+^ megakaryocytes from BM of PBS/E-Mac-injected site (tibia) and non-injected site (femur) were analyzed for DNA content. Plots from representative PBS-treated and E-Mac-treated leukemic mice four weeks post PBS/E-Mac treatment are shown. Percentages of mature megakaryocytes with 8N and greater ploidy (≥8N) are shown. **(G)** Statistical analysis of the percentages of mature megakaryocytes (≥8N) of CD41^+^ BM megakaryocytes. Data are analyzed by one-way ANOVA test. ***p < 0.001. Data are represented as mean ± SD (n = 4 mice for each group).

### MSCs reprogram macrophages and reduce leukemia burden through IL-6

Residual MSCs in leukemia-bearing mice lost the ability to secrete IL-6 (Figure 1G-H). IL-6 is critical for maintaining the stemness and function of MSCs^60^. The recovered host MSCs after donor MSC-treatment expressed comparable *Il6* mRNA as healthy MSCs (Figure 4E-G). Consistently, BM plasma IL-6 levels in MSC-treated leukemia-bearing mice were significantly elevated (> 3 folds) and were comparable to those in healthy mice (p < 0.001) (Figure 7A). To investigate the biological consequence of reduced IL-6 on MSCs, we analyzed the MSCs in *Il6*^-/-^ mice by *in vitro* proliferation assay. We observed that the *Il6*^-/-^ MSCs proliferated at a significantly slower speed than their WT counterparts *in vitro* (p < 0.001) (Figure. S18A). To investigate the role of IL-6 in MSC-mediated therapeutic effects, we directly injected IL-6 proteins (40 μg/kg) into leukemia-bearing mice. However, systemic IL-6 transfusion failed to suppress leukemia (Figure S18B), which indicates that MSC-educated leukemic macrophages are distinct from the conventional IL-6-dependent myeloid-derived immune-suppressor cells^61^. We speculated that IL-6 might function as an autocrine factor for MSCs to reprogram macrophages in leukemia-bearing mice. *In vitro* co-culture assay showed that *Il6*^-/-^ MSCs compromised their ability to induce *Arg1* expression in macrophages derived from leukemia-bearing mice (Figure 7B). In addition, intra-BM injection of *Il6*^-/-^ MSCs neither improved thrombopoiesis nor reduced tumor burden in leukemia-bearing mice (Figure 7C-E). Furthermore, loss of *Il6* significantly compromised MSCs’ ability to reprogram leukemic macrophages, demonstrated by mild increase of the ratios of Arg1^high^ macrophage subpopulation in *Il6*^-/-^ MSC-treated leukemic mice *in vivo* (Figure 7F-H). Taken together, these results demonstrate that autocrine IL-6 is essential for MSC-mediated reprogramming of macrophages and reduction of tumor burden in leukemia-bearing mice.

**Figure 7:**
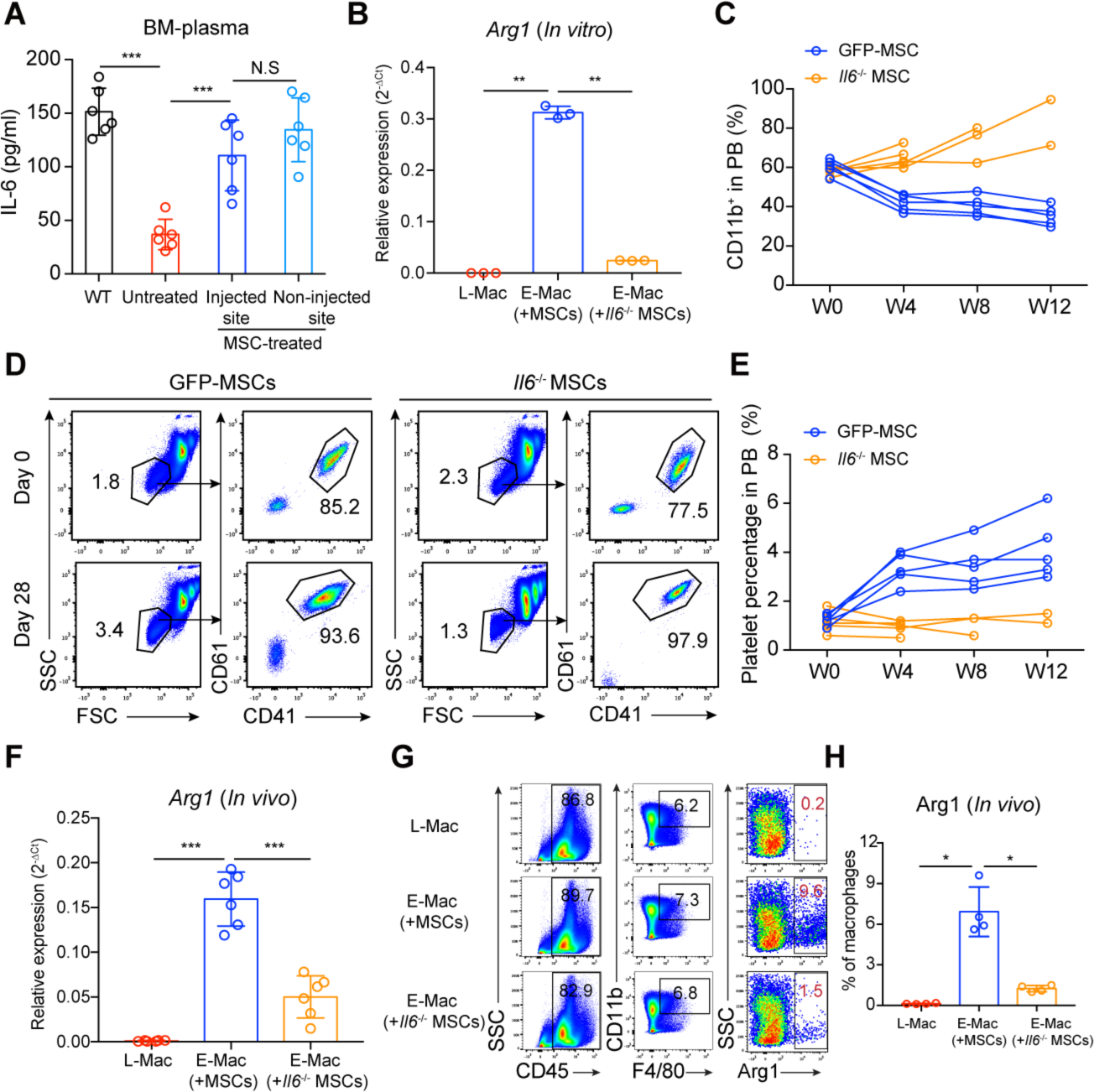
*Il6*^-/-^ MSCs neither reprogram macrophages nor suppress leukemia. **(A)** ELISA of IL-6 levels in BM plasma. The BM plasma from WT mice, untreated leukemia-bearing mice, and MSC-injected sites (tibias) and non-injected sites (femurs) of MSC-treated leukemia-bearing mice eight weeks post MSC treatment were analyzed. The BMNC of each tibia or femur were flushed out using 1 ml PBS, then the supernatants of each sample were collected for ELISA. Data are analyzed by one-way ANOVA test. ***p < 0.001. Data are represented as mean ± SD (n = 6 mice). **(B)** Q-PCR of *Arg1* expression levels in macrophages from leukemia-bearing mice after co-culture with different sources of MSCs *in vitro*. Macrophages from leukemia-bearing mice were co-cultured with MSCs from leukemia-bearing mice, *Il6*^-/-^ MSCs or GFP-MSCs *in vitro* for 12 h, separately. Then, one hundred thousand E-Mac (CD11b^+^F4/80^+^) were sorted and were analyzed the expression of *Arg1*. Y-axis shows the relative expression of *Arg1* in each group. Gapdh was used as a reference. Data are analyzed by one-way ANOVA test. **p < 0.01. Data are represented as mean ± SD (n = 3 repeats for each group). **(C)** Kinetic analysis of tumor burden (CD11b^+^) in PB of leukemia-bearing mice treated with GFP-MSCs or *Il6*^-/-^ MSCs (n = 5 mice for each group). **(D)** Flow cytometry analysis of platelets in *Il6*^-/-^ MSC-treated leukemia-bearing mice. Representative dot plots of platelet populations and quantitative gating, as identified by CD41 and CD61 staining in PB of GFP-MSC- or *Il6*^-/-^ MSC-treated leukemia-bearing mice. PB of leukemia-bearing mice was analyzed at week-0 and week-4 post-treatment with GFP-MSCs or *Il6*^-/-^ MSCs. **(E)** Kinetic analysis of platelets (PLT) in PB of leukemia-bearing mice treated with GFP-MSCs or *Il6*^-/-^ MSCs (n = 5 mice for each group). **(F)** Q-PCR of *Arg1* expression levels in macrophages isolated from *Il6*^-/-^ MSC-treated leukemia-bearing mice. One hundred thousand leukemic macrophages (CD11b^+^F4/80^+^) were sorted from the MSC-injected site (MSC-treated tibia) and non-injected site (femur) of leukemia-bearing mice 12h post-treatment. Y-axis shows the relative expression of *Arg1* in each group. Gapdh was used as reference. Data are analyzed by one-way ANOVA test. ***p < 0.001. Data are represented as mean ± SD (n = 6 mice for each group). **(G)** Representative intracellular staining plots of Arg1 proteins in MSC-reprogrammed leukemic macrophages (E-Mac, MSC-injected sites) and leukemic macrophages (L-Mac, Non-injected site) from GFP-MSC- or *Il6*^-/-^ MSC-treated leukemic mouse analyzed by flow cytometry. The ratios of Arg1^high^ macrophage subpopulation are shown from L-Mac and E-Mac. The bone marrow nucleated cells from leukemia-bearing mouse 12 h post-treatment were prepared for intracellular flow cytometry staining of Arg1. Macrophages are identified as CD45^+^CD11b^+^F4/80^+^. **(H)** Statistical analysis of the ratios of Arg1^high^ macrophage subpopulation. Data are analyzed by one-way ANOVA test. *p < 0.05. Data are represented as mean ± SD (n = 4 mice for each group).

## Discussion

Deteriorating BM microenvironment accompanies chronic leukemia progression. Here we unravel a *de novo* approach of reverting the impaired BM microenvironment by intra-BM injection of donor MSCs. Upon injection, the donor MSCs quickly reprogrammed local host BM macrophages to repair the niche, thus improving normal hematopoiesis and suppressing leukemia development. These effects of donor MSCs depend on the autocrine production of IL-6. Our studies reveal de novo mechanisms underlying MSC-mediated local BM microenvironment restoration that systemically suppress leukemia development.

Given the short-term lifespan of the exogenous MSC *in vivo*, it is surprising that local injection of donor MSCs results in long-term improvement of thrombopoiesis and reduction of tumor burden. Following injection, exogenous donor MSCs immediately reprogram host resident macrophages that further organize the overhaul of local BM microenvironment, including restoring the functions of host MSCs. There is a lot of evidence supporting the pivotal roles of macrophages in tissue repair^62, 63^. Donor MSCs can transiently release a key wave of tissue-repair factors, such as IL-6^45^, CCL7^50^, and CXCL12^47^, and reprogram host macrophages, subsequently resulting in the recovery of host MSCs. Recovered host MSCs further secreted much higher level of CCL2, CCL7, and CXCL12 that can further facilitate BM niche repair. Donor MSCs could also directly modulate the other niche cells, in addition to macrophages, to participate in BM niche repair^12^. Consequently, the restored local BM microenvironment outputs abundant hematopoiesis-improving cytokines, including IL-6^56^, and reduces tumor-growth-stimulating cytokines, such as GM-CSF^27, 55^. Thus, despite the short life-span, donor MSCs provide long-term thrombopoiesis improvement and tumor burden reduction through the stepwise microenvironment restoration.

Of note, IL-6 deficiency in MSCs markedly compromised the abilities of these cells to reprogram host resident macrophage. Transwell co-culture study has shown that direct interaction between MSCs and macrophages is a necessity for reprogramming host macrophages. Both autocrine production of IL-6 and reprogramming host macrophages by MSCs are required for MSC-mediated microenvironment restoration and leukemia inhibition. However, it is not clear how autocrine IL-6 controls the function of MSCs and how MSCs directly reprogram macrophages. The underlying molecular mechanisms warrant further investigation. Besides IL-6, it deserves additional investigations of whether other key molecules produced by MSCs are involved in the MSC-mediated microenvironment restoration and leukemia inhibition, such as nitric oxide^64^.

MSC treatment inhibits leukemia development in the *Nras* mutation-induced MPN/MDS-like disease model. We also attempted to broaden MSC treatment for acute leukemia in the MLL-AF9-initiated model (Figure S19A), in which impaired MSCs results in the reduction of osteogenesis and CXCL12 production^65^. Despite a mild elevation of platelet level, the intra-BM transfusion of donor MSCs failed to significantly improve normal hematopoiesis or suppress acute leukemia development (Figure S19B-H). Therefore, the intra-BM MSC treatment might be beneficial for MPN/MDS leukemias, such as JMML and CMML, but insufficient for suppressing acute leukemia. Although MSC application could be an effective therapeutic regimen for patients with MPN/MDS-subtype leukemia, combination therapy of conventional approaches with local MSC transfusion might be required to achieve therapeutic outcomes for acute leukemia with impaired BM microenvironment.

## Acknowledgments

This work was supported by grants from the Strategic Priority Research Program of Chinese Academy of Sciences (XDA16010601), the Chinese Ministry of Science and Technology (2015CB964401, 2016YFA0100601, 2016YFA0100600, 2017YFA0103401, and 2015CB964902), the Major Research and Development Project of Guangzhou Regenerative Medicine and Health Guangdong Laboratory (2018GZR110104006, 2018GZR0201008), the CAS Key Research Program of Frontier Sciences (QYZDB-SSW-SMC057), the Health and Medical Care Collaborative Innovation Program of Guangzhou Scientific and Technology (201803040017), CAMS Innovation Fund for Medical Sciences (2016-12M-1-002), the General Program from Guangzhou Scientific and Technological Project (201707010157), the Science and Technology Planning Project of Guangdong Province (2017B030314056, 2017B020230004), the grants from the National Natural Science Foundation of China (Grant No 31471117, 31271457, 81470281, 81421002, 81730006, 31600948, and 81861148029), the CAMS Initiative for Innovative Medicine (2016-I2M-1-017) and the grants from NIH, USA (AI079087, D.W. and HL130724 D.W.)

## Author Contributions

C.X.X. performed research, analyzed data and wrote the manuscript; D.Y. and Q.T.W. analyzed RNA-Seq data; T.J.W., H.C., P.Q.Z., K.T.W., X.F.L., Y.G., S.H.M., L.X. and Y.X.G. performed experiments; S.H., J.D., X.D., Y.Q.L., X.F.Z., Y.F.S. and S.X. discussed the manuscript; D.W. discussed the project and wrote the manuscript. T.C. and J.Y.W. designed the research and wrote the manuscript.

## References

1. Kfoury Y, Scadden DT. Mesenchymal cell contributions to the stem cell niche. Cell Stem Cell. 2015;16(3):239–253.

2. Lim M, Pang Y, Ma S, et al. Altered mesenchymal niche cells impede generation of normal hematopoietic progenitor cells in leukemic bone marrow. Leukemia. 2016;30(1):154–162.

3. Raaijmakers MH, Mukherjee S, Guo S, et al. Bone progenitor dysfunction induces myelodysplasia and secondary leukaemia. Nature. 2010;464(7290):852–857.

4. Schepers K, Pietras EM, Reynaud D, et al. Myeloproliferative neoplasia remodels the endosteal bone marrow niche into a self-reinforcing leukemic niche. Cell Stem Cell. 2013;13(3):285–299.

5. Bowers M, Zhang B, Ho Y, Agarwal P, Chen CC, Bhatia R. Osteoblast ablation reduces normal long-term hematopoietic stem cell self-renewal but accelerates leukemia development. Blood. 2015;125(17):2678–2688.

6. Dong L, Yu WM, Zheng H, et al. Leukaemogenic effects of Ptpn11 activating mutations in the stem cell microenvironment. Nature. 2016;539(7628):304–308.

7. Blau O, Baldus CD, Hofmann WK, et al. Mesenchymal stromal cells of myelodysplastic syndrome and acute myeloid leukemia patients have distinct genetic abnormalities compared with leukemic blasts. Blood. 2011;118(20):5583–5592.

8. Pandis N, Bardi G, Sfikas K, Panayotopoulos N, Tserkezoglou A, Fotiou S. Complex chromosome rearrangements involving 12q14 in two uterine leiomyomas. Cancer Genet Cytogenet. 1990;49(1):51–56.

9. Medyouf H, Mossner M, Jann JC, et al. Myelodysplastic cells in patients reprogram mesenchymal stromal cells to establish a transplantable stem cell niche disease unit. Cell Stem Cell. 2014;14(6):824–837.

10. Ren G, Zhang L, Zhao X, et al. Mesenchymal stem cell-mediated immunosuppression occurs via concerted action of chemokines and nitric oxide. Cell Stem Cell. 2008;2(2):141–150.

11. Prockop DJ. Inflammation, fibrosis, and modulation of the process by mesenchymal stem/stromal cells. Matrix Biol. 2016;51:7–13.

12. Shi Y, Wang Y, Li Q, et al. Immunoregulatory mechanisms of mesenchymal stem and stromal cells in inflammatory diseases. Nat Rev Nephrol. 2018;14(8):493–507.

13. Le Blanc K, Mougiakakos D. Multipotent mesenchymal stromal cells and the innate immune system. Nat Rev Immunol. 2012;12(5):383–396.

14. Mittal M, Tiruppathi C, Nepal S, et al. TNFalpha-stimulated gene-6 (TSG6) activates macrophage phenotype transition to prevent inflammatory lung injury. Proc Natl Acad Sci U S A. 2016;113(50):E8151–E8158.

15. Wang G, Cao K, Liu K, et al. Kynurenic acid, an IDO metabolite, controls TSG-6-mediated immunosuppression of human mesenchymal stem cells. Cell Death Differ. 2018;25(7):1209–1223.

16. Du L, Lin L, Li Q, et al. IGF-2 Preprograms Maturing Macrophages to Acquire Oxidative Phosphorylation-Dependent Anti-inflammatory Properties. Cell Metab. 2019.

17. Eggenhofer E, Benseler V, Kroemer A, et al. Mesenchymal stem cells are short-lived and do not migrate beyond the lungs after intravenous infusion. Front Immunol. 2012;3:297.

18. Ehninger A, Trumpp A. The bone marrow stem cell niche grows up: mesenchymal stem cells and macrophages move in. J Exp Med. 2011;208(3):421–428.

19. Chen J, Yao Y, Gong C, et al. CCL18 from tumor-associated macrophages promotes breast cancer metastasis via PITPNM3. Cancer Cell. 2011;19(4):541–555.

20. Gubin MM, Esaulova E, Ward JP, et al. High-Dimensional Analysis Delineates Myeloid and Lymphoid Compartment Remodeling during Successful Immune-Checkpoint Cancer Therapy. Cell. 2018;175(4):1014–1030 e1019.

21. Chen CC, Wang L, Plikus MV, et al. Organ-level quorum sensing directs regeneration in hair stem cell populations. Cell. 2015;161(2):277–290.

22. Liu C, Wu C, Yang Q, et al. Macrophages Mediate the Repair of Brain Vascular Rupture through Direct Physical Adhesion and Mechanical Traction. Immunity. 2016;44(5):1162–1176.

23. Winkler IG, Sims NA, Pettit AR, et al. Bone marrow macrophages maintain hematopoietic stem cell (HSC) niches and their depletion mobilizes HSCs. Blood. 2010;116(23):4815–4828.

24. Bosurgi L, Cao YG, Cabeza-Cabrerizo M, et al. Macrophage function in tissue repair and remodeling requires IL-4 or IL-13 with apoptotic cells. Science. 2017;356(6342):1072–1076.

25. Cho DI, Kim MR, Jeong HY, et al. Mesenchymal stem cells reciprocally regulate the M1/M2 balance in mouse bone marrow-derived macrophages. Exp Mol Med. 2014;46:e70.

26. Nemeth K, Leelahavanichkul A, Yuen PS, et al. Bone marrow stromal cells attenuate sepsis via prostaglandin E(2)-dependent reprogramming of host macrophages to increase their interleukin-10 production. Nat Med. 2009;15(1):42–49.

27. Wang J, Liu Y, Li Z, et al. Endogenous oncogenic Nras mutation promotes aberrant GM-CSF signaling in granulocytic/monocytic precursors in a murine model of chronic myelomonocytic leukemia. Blood. 2010;116(26):5991–6002.

28. Wang J, Liu Y, Li Z, et al. Endogenous oncogenic Nras mutation initiates hematopoietic malignancies in a dose- and cell type-dependent manner. Blood. 2011;118(2):368–379.

29. Wang JY, Kong GY, Liu YG, et al. Nras(G12D/+) promotes leukemogenesis by aberrantly regulating hematopoietic stem cell functions. Blood. 2013;121(26):5203–5207.

30. Tang F, Barbacioru C, Nordman E, et al. RNA-Seq analysis to capture the transcriptome landscape of a single cell. Nat Protoc. 2010;5(3):516–535.

31. Kim D, Langmead B, Salzberg SL. HISAT: a fast spliced aligner with low memory requirements. Nat Methods. 2015;12(4):357–360.

32. Liao Y, Smyth GK, Shi W. featureCounts: an efficient general purpose program for assigning sequence reads to genomic features. Bioinformatics. 2014;30(7):923–930.

33. Love MI, Huber W, Anders S. Moderated estimation of fold change and dispersion for RNA-seq data with DESeq2. Genome Biol. 2014;15(12):550.

34. Subramanian A, Tamayo P, Mootha VK, et al. Gene set enrichment analysis: a knowledge-based approach for interpreting genome-wide expression profiles. Proc Natl Acad Sci U S A. 2005;102(43):15545–15550.

35. Yu G, Wang LG, Han Y, He QY. clusterProfiler: an R package for comparing biological themes among gene clusters. OMICS. 2012;16(5):284–287.

36. Freeman BT, Jung JP, Ogle BM. Single-Cell RNA-Seq of Bone Marrow-Derived Mesenchymal Stem Cells Reveals Unique Profiles of Lineage Priming. PLoS One. 2015;10(9):e0136199.

37. Song L, Webb NE, Song Y, Tuan RS. Identification and functional analysis of candidate genes regulating mesenchymal stem cell self-renewal and multipotency. Stem Cells. 2006;24(7):1707–1718.

38. Rostovskaya M, Anastassiadis K. Differential expression of surface markers in mouse bone marrow mesenchymal stromal cell subpopulations with distinct lineage commitment. PLoS One. 2012;7(12):e51221.

39. Delorme B, Ringe J, Pontikoglou C, et al. Specific lineage-priming of bone marrow mesenchymal stem cells provides the molecular framework for their plasticity. Stem Cells. 2009;27(5):1142–1151.

40. Medina RJ, O’Neill CL, O’Doherty TM, et al. Myeloid angiogenic cells act as alternative M2 macrophages and modulate angiogenesis through interleukin-8. Mol Med. 2011;17(9-10):1045–1055.

41. Arango Duque G, Descoteaux A. Macrophage cytokines: involvement in immunity and infectious diseases. Front Immunol. 2014;5:491.

42. Cavaillon JM. Cytokines and macrophages. Biomed Pharmacother. 1994;48(10):445–453.

43. Li Q, Haigis KM, McDaniel A, et al. Hematopoiesis and leukemogenesis in mice expressing oncogenic NrasG12D from the endogenous locus. Blood. 2011;117(6):2022–2032.

44. Gnecchi M, Danieli P, Malpasso G, Ciuffreda MC. Paracrine Mechanisms of Mesenchymal Stem Cells in Tissue Repair. Methods Mol Biol. 2016;1416:123–146.

45. Gallucci RM, Simeonova PP, Matheson JM, et al. Impaired cutaneous wound healing in interleukin-6-deficient and immunosuppressed mice. FASEB J. 2000;14(15):2525–2531.

46. Castela M, Nassar D, Sbeih M, Jachiet M, Wang Z, Aractingi S. Ccl2/Ccr2 signalling recruits a distinct fetal microchimeric population that rescues delayed maternal wound healing. Nat Commun. 2017;8:15463.

47. Kato T, Khanh VC, Sato K, et al. SDF-1 improves wound healing ability of glucocorticoid-treated adipose tissue-derived mesenchymal stem cells. Biochem Biophys Res Commun. 2017;493(2):1010–1017.

48. Hayashi Y, Murakami M, Kawamura R, Ishizaka R, Fukuta O, Nakashima M. CXCL14 and MCP1 are potent trophic factors associated with cell migration and angiogenesis leading to higher regenerative potential of dental pulp side population cells. Stem Cell Res Ther. 2015;6:111.

49. Sims NA, Jenkins BJ, Nakamura A, et al. Interleukin-11 receptor signaling is required for normal bone remodeling. J Bone Miner Res. 2005;20(7):1093–1102.

50. Schenk S, Mal N, Finan A, et al. Monocyte chemotactic protein-3 is a myocardial mesenchymal stem cell homing factor. Stem Cells. 2007;25(1):245–251.

51. Kleppe M, Kwak M, Koppikar P, et al. JAK-STAT pathway activation in malignant and nonmalignant cells contributes to MPN pathogenesis and therapeutic response. Cancer Discov. 2015;5(3):316–331.

52. Pang L, Weiss MJ, Poncz M. Megakaryocyte biology and related disorders. J Clin Invest. 2005;115(12):3332–3338.

53. Zhu H, Guo ZK, Jiang XX, et al. A protocol for isolation and culture of mesenchymal stem cells from mouse compact bone. Nat Protoc. 2010;5(3):550–560.

54. Rombouts WJ, Ploemacher RE. Primary murine MSC show highly efficient homing to the bone marrow but lose homing ability following culture. Leukemia. 2003;17(1):160–170.

55. Padron E, Painter JS, Kunigal S, et al. GM-CSF-dependent pSTAT5 sensitivity is a feature with therapeutic potential in chronic myelomonocytic leukemia. Blood. 2013;121(25):5068–5077.

56. Kaser A, Brandacher G, Steurer W, et al. Interleukin-6 stimulates thrombopoiesis through thrombopoietin: role in inflammatory thrombocytosis. Blood. 2001;98(9):2720–2725.

57. de Lima M, McNiece I, Robinson SN, et al. Cord-blood engraftment with ex vivo mesenchymal-cell coculture. N Engl J Med. 2012;367(24):2305–2315.

58. Ohnishi H, Kobayashi H, Okazawa H, et al. Ectodomain shedding of SHPS-1 and its role in regulation of cell migration. J Biol Chem. 2004;279(27):27878–27887.

59. Hocking AM. The Role of Chemokines in Mesenchymal Stem Cell Homing to Wounds. Adv Wound Care (New Rochelle). 2015;4(11):623–630.

60. Pricola KL, Kuhn NZ, Haleem-Smith H, Song Y, Tuan RS. Interleukin-6 maintains bone marrow-derived mesenchymal stem cell stemness by an ERK1/2-dependent mechanism. J Cell Biochem. 2009;108(3):577–588.

61. Marigo I, Bosio E, Solito S, et al. Tumor-induced tolerance and immune suppression depend on the C/EBPbeta transcription factor. Immunity. 2010;32(6):790–802.

62. Wynn TA, Vannella KM. Macrophages in Tissue Repair, Regeneration, and Fibrosis. Immunity. 2016;44(3):450–462.

63. Liu C, Wu CA, Yang QF, et al. Macrophages Mediate the Repair of Brain Vascular Rupture through Direct Physical Adhesion and Mechanical Traction. Immunity. 2016;44(5):1162–1176.

64. Trento C, Marigo I, Pievani A, et al. Bone marrow mesenchymal stromal cells induce nitric oxide synthase-dependent differentiation of CD11b(+) cells that expedite hematopoietic recovery. Haematologica. 2017;102(5):818–825.

65. Hanoun M, Zhang D, Mizoguchi T, et al. Acute myelogenous leukemia-induced sympathetic neuropathy promotes malignancy in an altered hematopoietic stem cell niche. Cell Stem Cell. 2014;15(3):365–375.

