## Supplementary information for "Mesenchymal stem cells suppress leukemia via macrophage-mediated functional restoration of bone marrow microenvironment"

[1]State Key Laboratory of Experimental Hematology, CAS Key Laboratory of Regenerative Biology, Guangzhou Institutes of Biomedicine and Health, Chinese Academy of Sciences, Guangzhou, China; [2]State Key Laboratory of Experimental Hematology & National Clinical Research Center for Blood Diseases, Institute of Hematology & Blood Diseases Hospital, Chinese Academy of Medical Sciences & Peking Union Medical College, Tianjin, China; [3]Guangzhou Regenerative Medicine and Health-Guangdong Laboratory (GRMH-GDL), Guangzhou, China; [4]Guangdong Provincial Key Laboratory of Stem cell and Regenerative Medicine, Guangzhou Institutes of Biomedicine and Health, Chinese Academy of Sciences, Guangzhou, China; [5]Center for Stem Cell Medicine & Department of Stem Cell and Regenerative Medicine, Chinese Academy of Medical Sciences & Peking Union Medical College, Tianjin, China; [6]Joint School of Life Sciences, Guangzhou Institutes of Biomedicine and Health, Guangzhou Medical University, Guangzhou, China; [7]University of Chinese Academy of Sciences, Beijing, China; [8]Department of Hematology, First Affiliated Hospital; Institute of Hematology, School of Medicine; Key Laboratory for Regenerative Medicine of Ministry of Education; Jinan University, Guangzhou, China; [9]Department of Hematology, Guangdong Provincial People's Hospital (Guangdong Academy of Medical Sciences), Guangzhou, Guangdong, China; [10]South China University of Technology, Guangzhou, China; [11]The First Affiliated Hospital of Soochow University, State Key Laboratory of Radiation Medicine and Protection, Institutes for Translational Medicine, Soochow University, Suzhou, China; [12]National Key Laboratory of Medical Immunology & Institute of Immunology, Second Military Medical University, Shanghai, China; [13]Blood Research Institute, Versiti, Milwaukee, WI, USA; [14]Biomedical Research Center of South China, College of Life Sciences, Fujian Normal University, Fuzhou, China.

[†]Equal contributors.

[*]Correspondences:

Jinyong Wang (J.W.),

Tao Cheng (T.C.),

### Supplementary materials and methods

**MLL-AF9 AML mouse model.** We used a non-irradiated acute myeloid leukemia mouse model described previously[1].

**Flow cytometry analysis.** Antibodies for hematopoietic lineage analysis: FITC-TER-119 (TER-119), PerCP-Cyanine5.5-CD45.2 (104), APC-Thy1.2 (53-2.1), APC-CD3e (145-2C11), PE-CD19 (1D3), PE-Cy7-CD11b (M1/70), APC-eFluor®780-Gr-1 (RB6-8C5), FITC-CD41 (MWReg30), PE-CD61 (2C9.G3), APC-eFluor®780-F4/80 (BM8), APC-CD206 (C068C2), and PE-CD11c (N418) antibodies were purchased from eBiosciences or Biolegend. DAPI was used to exclude dead cells.

For MSC analysis, BMNC were stained with the following antibodies: APC-Ter119 (TER-119), APC-CD45 (30-F11), PE-Cy7-CD31 (WM-59), APC-eFluor®780-Sca1 (D7), PE-CD51 (RMV-7), and PerCP-Cyanine5.5-CD146 (ME-9F1) were purchased from eBiosciences or Biolegend. DAPI was used to exclude dead cells.

For myeloid progenitors staining, BM cells were stained with the following antibodies: CD2 (RM2-5), CD3e (145-2C11), CD4 (RM4-5), CD8a (53-6.7), Ter119 (TER-119), CD11b (M1/70), B220 (6B2), Gr1 (RB6-8C5), IL-7R (A7R34), Sca1 (E13-161.7), c-kit (2B8), CD34 (RAM34), and CD16/32 (93). DAPI was used to exclude dead cells.

For megakaryocyte maturation detection, BM cells were stained with labeled with CD41-FITC (MWReg30). Then cells were fixed using cold 70% ethanol. After washing, the fixed cells were resuspended in propidium iodide.

For platelet staining and counting, 5 μL fresh whole blood was collected. Whole blood sample was blocked with anti-mouse CD16/32, then was stained with anti-mouse CD41-FITC and anti-mouse CD61-PE at room temperature for 20 mins. Then 1 mL of cold 1% PFA solution and 50 μL absolute counting beads (C36950, Invitrogen) were added to each sample. The sample was fixed on ice for at least 30 mins.

For intracellular flow cytometry staining, BM cells were blocked with anti-mouse CD16/32 and stained with surface markers for 20 min on ice. Then cells were fixed using IC fixation buffer (88-8824-00, eBiosciences) and pulsed vortex to mix. After washing, cells were resuspended with 1X permeabilization buffer and stained with the intracellular antibodies for 40 min at room temperature. Finally, the cells were resuspended with FCS buffer for analysis. The following intracellular antibodies were used: FITC-CXCL-12 (79018, Invitrogen), APC-Arginase 1 (A1exF5, eBiosciences), PE/Cy7-GM-CSF (MP1-22E9, Biolegend).

The stained cells were analyzed on LSR Fortessa (BD Bioscience), then the data were analyzed using Flowjo software (FlowJo).

**Preparation of BMNC.** Mice were sacrificed, and BM cells were isolated by flushing out the tibias and femurs using DPBS containing 2% FBS. The compact bones were dissected into ~2 mm fragments and transferred with 5ml of 1 mg/ml collagenase II solution into a 50 ml tube. The tubes were incubated in a shaker (< 110 rpm) at $37^{o}C$ for 1-2 hours. BMNC from BM cells and compact bones were mixed and filtered through a 70 μm cell strainer (BD Falcon) to obtain a single-cell suspension.

**MSC sorting.** BMNC mixtures of BM and compact bones from the control mice (age-matched wild-type, CD11b$^{+}$% in PB = 10-15%), leukemia-bearing mice (CD11b$^{+}$% in PB = 35%-45%) and leukemia-bearing mice eight weeks post-treatment with GFP$^{+}$ MSCs were isolated as previously described. After lysis of red blood cells, BMNC were blocked by Fc blocker, and incubated with biotin-conjugated anti-CD45 antibody and enriched by streptavidin magnetic beads (Miltenyi Bitec). The enriched CD45$^{-}$ cells were stained with the following antibodies: APC-Ter119 (TER-

119), APC-CD45 (30-F11), streptavidin-APC, PE-Cy7-CD31 (WM-59), APC-eFluor®780-Sca1 (D7), PE-CD51 (RMV-7) were purchased from eBiosciences. DAPI was used to exclude dead cells. MSCs were sorted by the gating strategy defining as $GFP^-Ter119^-CD45^-CD31^-Sca1^+CD51^+$ using AriaII (BD Bioscience) and subsequently prepared for RNA-Seq.

**Mouse colony-forming unit-fibroblast (CFU-F) assay.** For analyzing the quantity of functional MSCs, BMNC equivalent to 100 MSCs from each mouse were used as cell input for individual wells (six-well plate). BMNC were suspended into 2 ml of mouse complete MesenCult™ medium (Catalog 05513, StemCell Technology), then seeded into the individual wells. BMNC were incubated at 37°C with 5% $CO_2$ in a humidified chamber. Half-medium change was performed on day 7. After 14 days, the wells were washed once with DPBS and fixed by ice-cold ethanol, and then stained with giemsa stain at RT. After washing, colonies with more than 20 spindle-shaped cells per colony were counted. Three replicates of each sample were performed.

**Isolation and expansion of mouse MSCs.** MSCs were isolated from cell mixture of compact bones and BM cells of 3-4 weeks old healthy GFP mice (n = 100) or $Il6^{-/-}$ mice, as previously reported with minor modifications[2]. Briefly, the BM cavities were flushed to thoroughly deplete hematopoietic cells. The compact bones were dissected into ~2 mm fragments and transferred with 5ml of 1 mg/ml collagenase II solution into a 50 ml tube. The tubes were incubated in a shaker (< 110 rpm) at 37°C for 1-2 hours. The fragments were washed three times and cultivated in complete MSC culture medium ( α-MEM (Gibco) supplemented with 10% FBS (Gibco) and 1% penicillin/streptomycin (Invitrogen)) in a 6 cm dish. Besides, MSCs from the BM cells were sorted ($Ter119^-CD45^-CD31^-Sca1^+CD51^+CD146^+$) directly into MSC culture medium. These two sources of MSCs from compact bones and BM cells were mixed for further isolation and expansion. The bone fragments were removed, and culture medium was replaced after three times' washing on the third day. After culture for five days, the adherent cells were harvested by 0.25% trypsin's digestion and passaged. The culture medium was changed every 48 hours and passaged at a split ratio of 1:3 every 3-4 days. The expanded MSCs (Passage 2) were cryopreserved with 90% DMSO and 10% FBS in liquid nitrogen for transfusion. The cryopreserved P2 MSCs were recovered and cultured for 4-5 days, phenotypically identified, and collected in DPBS (2.5 × 10^7/ml) for transfusion.

**Osteogenic differentiation of MSCs.** Healthy MSCs or leukemic MSCs were plated in MSC growth medium at a density of 2 × 10^4 cells/cm2 in 6-well plates pre-coated with gelatin solution and incubated at $37^{o}$C under 5% CO2 in a humidified incubator. When cells were approximately 60-70% confluent, the growth medium was removed and 2 ml mouse mesenchymal stem cell osteogenic differentiation medium (MUBMX-90021, Cyagen) was added. The medium was replaced every three days. After 3 weeks' induction, cells were fixed with 4% neutral formaldehyde and stained with alizarin red. Images were captured and mineralization area was calculated by Image-Pro software.

**Adipogenic differentiation of MSCs.** Healthy MSCs or leukemic MSCs were plated in MSC growth medium at a density of 2 × 10^4 cells/cm2 in 6-well plates pre-coated with gelatin solution, and incubated at $37^{o}$C under 5% CO2 in a humidified incubator. When cells were approximately 100% confluent, the growth medium was removed and 2 ml mouse mesenchymal stem cell adipogenic differentiation induction medium A (MUBMD-90031, Cyagen) was added. Three days later, change the medium to maintenance medium B by replacing the induction medium A. 24 hours later, change the medium back to induction medium A. After 3-5 cycles' induction, cells were fixed with 4% neutral formaldehyde and stained with oil red O. Images were captured and lipid droplets area was calculated by Image-Pro software.

**Complete blood count (CBC).** For mouse samples, 100 μl PB from each mouse was collected into 1.5 ml anticoagulation tube and diluted with the same volume of PBS, then performed complete blood count by automatic blood analyzer (Abbott, CD3700SL).

**Enzyme-linked immunosorbent assay (ELISA) assay.** For mouse samples, serum was collected from PB of control mice and leukemia-bearing mice; and BM plasma samples were collected from tibias and femurs of control mice and leukemia-bearing mice by flushing out the BM using 1-2 ml PBS. The supernatants were collected for ELISA after centrifugation. The concentration of cytokines, mouse IL-6 (CME0006-96) and mouse GM-CSF (CME0026-96), were measured using ELISA kits (Beijing 4A Biotech Corporation, China) according to the manufacture's instruction.

**Hematoxylin and eosin-staining and immunohistochemistry.** Tibias from control mice and leukemic mice were fixed in 4% paraformaldehyde, following dehydration by 30% sucrose, and embedded in paraffin for sectioning. Then tibias were processed for histologic hematoxylin and eosin-staining (Pathology lab, GIBH). For CD11b staining, tibia sections were processed to quench endogenous peroxidase activity and blocked by blocking buffer (5% normal goat serum and 0.3% Triton X-100 in PBS). Sections were incubated with the primary anti-CD11b polyclonal antibody (ab133357, Abcam, 1:1000) overnight at 4$^{o}$C, followed by secondary biotinylated anti-rabbit antibody (goat anti-rabbit IgG-HRP, sc-2004, Santa Cruz, 1:1000 dilution) for 1 hour at room temperature. A mixture of DAB substrate and DAB chromogen was used for antigen visualization. Sections were counterstained with hematoxylin and mounted in neutral balsam. Images were captured and processed using Motic digital slice scan system (VMV1 VMDPCS, Motic) and DSS scanner software.

**Indirect immunofluorescence assay (IFA).**

Sorted MSCs or macrophages (100 cells in 5μl) were directly pipetted onto the microscope slides (Thermo Scientific™ Superfrost Plus™ and Colorfrost Plus™ adhesion slides, 4951PLUS) and incubated at room temperature for 10 min. Upon the solution was completely dry, the cells were fixed with 4%PFA for 10 min following with 0.15%Triton X-100 permeabilization for 2 min at room temperature. To avoid non-specific antibody binding, the cells were blocked in 1% BSA/PBS for 1-2h at room temperature and then incubated with the primary antibody in 1% BSA in PBS overnight at 4℃. The following primary antibodies were used: APC-IL-6 (MP5-20F3, Biolegend), APC-MCP-1 (2H5, Biolegend), APC-TNF-α (MP6-XT22, Biolegend), PE-CCR7 (4B12, Biolegend), IL-1α rabbit polyclonal antibody (16765-1-AP, proteintech). Slides were washed three times in PBS and incubated with corresponding secondary antibodies for 1h at room temperature in 1% BSA in PBS. The following secondary antibodies were used: donkey anti-rat Alexa Fluor® 647 (ab150155, Abcam), donkey anti-rabbit Alexa Fluor® 555 (ab150074, Abcam). After washing the slides, the cells were incubated with DAPI solution for 10 min. Confocal analysis was performed at high resolution with a Zeiss laser scanning confocal microscope, LSM-800. The images were processed with ZEN 2012 software (blue edition).

**mIL-6 treatment for leukemia-bearing mice.** A dose of 40 μg/kg mIL-6 was injected into each leukemia-bearing mouse by intraperitoneal injection. mIL-6 was injected once every day and continued in a time window of 4 weeks. Analysis of tumor burden ($CD11b^+$ cells) in PB was performed every two weeks.

**Quantitative real-time PCR.** For analysis of mRNA expression levels of related genes in MSCs and $CD11b^+$ leukemic cells from the co-culture assay, 1 × 10^5 target cells of each sample were sorted by flow cytometry using Aria II. Total RNA was extracted using the RNeasy Micro Kit (Cat NO. 74004, QIAGEN). On-column DNase digestion of the samples was performed following the manufacturer's instruction.  First strand cDNA was synthesized from 100 ng of total RNA in 20 μl final volume, using the ReverTra Ace qPCR RT Master Mix kit (FSQ-301, TOYOBO) according to the manufacturer's instructions. Real-time quantitative PCR assays were carried out in a BioRad CFX96 Real-Time PCR Detection System instrument (Bio-Rad) using standard PCR conditions. Triplicates of all reactions were performed. GAPDH gene was used as a reference for differential expression comparison. The primer sequences of all related genes are shown in Table S1.

**Table S1. Real-time quantitative PCR primer sequences.**

| Gene | Forward (5'-3') | Reverse (5'-3') |
|---|---|---|
| *Gapdh* | TGGTGAAGGTCGGTGTGAACG | CAATGAAGGGGTCGTTGATGGC |
| *Il6* | TAGTCCTTCCTACCCCAATTTCC | TTGGTCCTTAGCCACTCCTTC |
| *Arg1* | CATTGGCTTGCGAGACGTAGAC | GCTGAAGGTCTCTTCCATCACC |

### Supplementary figures and figure legends

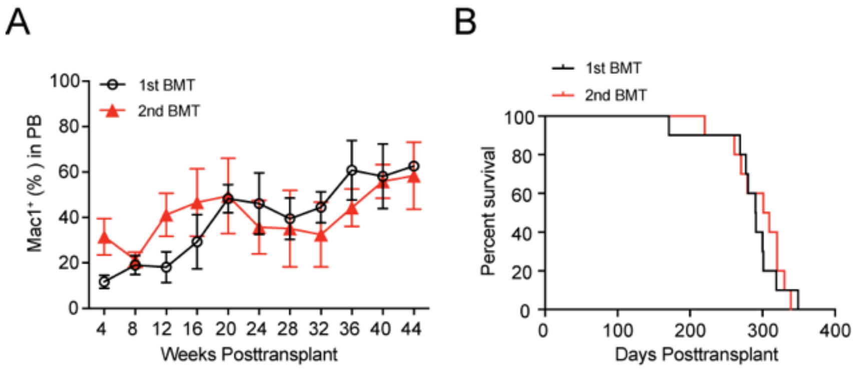


**Figure S1: Kinetics of tumor burden (CD11b$^+$) and survival curves of primary and secondary leukemia-bearing mice**

For the primary transplantation, 0.3 million sorted CD45.2$^+$ BMNC from *LSL Nras/+*; *Vav-Cre* mice were transplanted into sublethally irradiated (6.5 Gy) individual recipients (CD45.1 strain). The leukemia-bearing mice (CD11b$^+$% of PB > 60%) were used as donors for secondary transplantation (2$^{nd}$ BMT) 36 weeks after primary transplantation. For secondary transplantation, one million sorted CD45.2$^+$ BMNC of leukemia-bearing mice were transplanted into sublethally irradiated (6.5 Gy) individual recipients (CD45.1 strain). **(A)** Kinetic analysis of donor-derived myeloid cells (CD11b$^+$) in PB of primary transplantation (1$^{st}$ BMT) and secondary transplantation (2$^{nd}$ BMT) recipients. Flow cytometry analysis of CD11b$^+$ cells in PB was performed monthly. Data are represented as mean ± SD (n = 10). **(B)** Kaplan-Meier survival of primary and secondary transplantation recipients. Kaplan-Meier survival curves of 1$^{st}$ BMT (black line, n = 10, Median survival = 290.5 days) and 2$^{nd}$ BMT (red line, n = 10, Median survival = 305 days) leukemia-bearing mice are shown. Log-rank (Mantel-Cox) test: p = 0.6105.

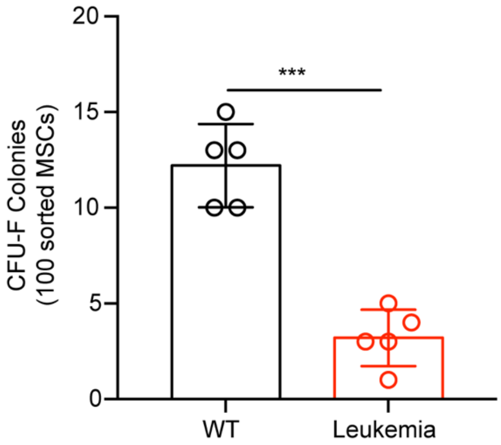


**Figure S2: Leukemic MSCs form fewer colonies than healthy counterparts *in vitro***

Statistical analysis of functional MSCs in CFU-F assay. One hundred primary MSCs were sorted from wild type mice or leukemia-bearing mice as cell input for CFU-F assay. Data are analyzed by unpaired student's t-test (two-tailed). ***$p < 0.001$. Five repeats were performed in three independent experiments. Data are represented as mean ± SD.

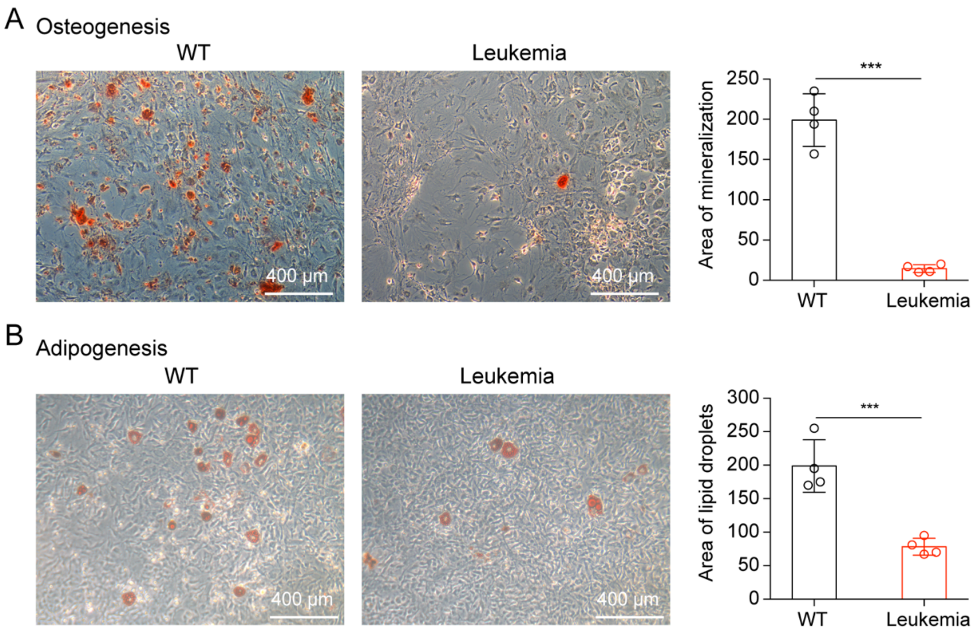


**Figure S3: Leukemic MSCs show reduced abilities of osteogenic and adipogenic differentiation *in vitro***

**(A)** Osteogenic differentiation of MSCs isolated from wild type mice and leukemia-bearing mice. MSCs isolated from WT mice or leukemia-bearing mice were cultured in osteogenic differentiation medium for three weeks. On day 21, the differentiated cells were fixed with 4% neutral formaldehyde and stained with alizarin red. Images were captured and the formed mineralization area was calculated by Image-Pro software. Bars = 400 μm. Data are analyzed by unpaired student's t-test (two-tailed). ***$P < 0.001$. Data are represented as mean ± SD (n = 4 replicates). **(B)** Adipogenic differentiation of MSCs isolated from wild type mice and leukemia-bearing mice. MSCs isolated from WT mice or leukemia-bearing mice were cultured in adipogenic differentiation medium for three weeks. On day 21, the differentiated cells were fixed with 4% neutral formaldehyde and stained with oil red O. Images were captured, and the formed lipid droplets area was calculated by Image-Pro software. Bars = 400 μm. Data are analyzed by unpaired student's t-test (two-tailed). ***$P < 0.001$. Data are represented as mean ± SD (n = 4 replicates).

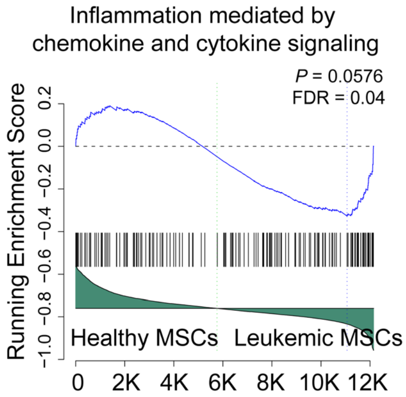


**Figure S4: GSEA of the inflammation mediated by chemokine and cytokine signaling in healthy MSCs and leukemic MSCs**

DESeq2 normalized values of the expression data were used for GSEA analysis.

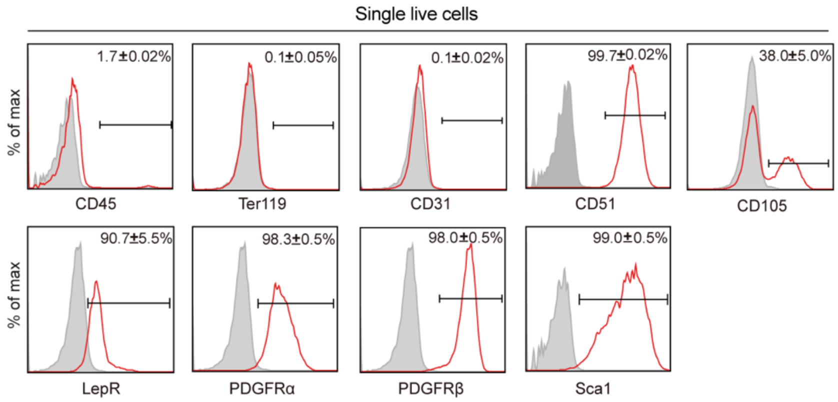


**Figure S5: Characterization of donor MSCs prior injection**

Phenotypic identification of the MSCs before injection. The expanded MSCs (passage 2) were analyzed by flow cytometry. For each batch of MSC preparation, MSCs were isolated from the compact bones and BMNC of twenty GFP mice (3–4 weeks old). The phenotypic identification of MSCs was performed before injection. MSCs are identified by the following markers: $CD45^-$, $Ter119^-$, $CD31^-$, $CD51^+$, $CD105^+$, $LepR^+$, PDGFR$\alpha^+$, PDGFR $\beta^+$, $Sca1^+$, as shown in the red histograms. The isotype controls of each antibody were used as negative controls, as shown in the grey histograms. Data are shown as mean ± SD, which were from three independent experiments.

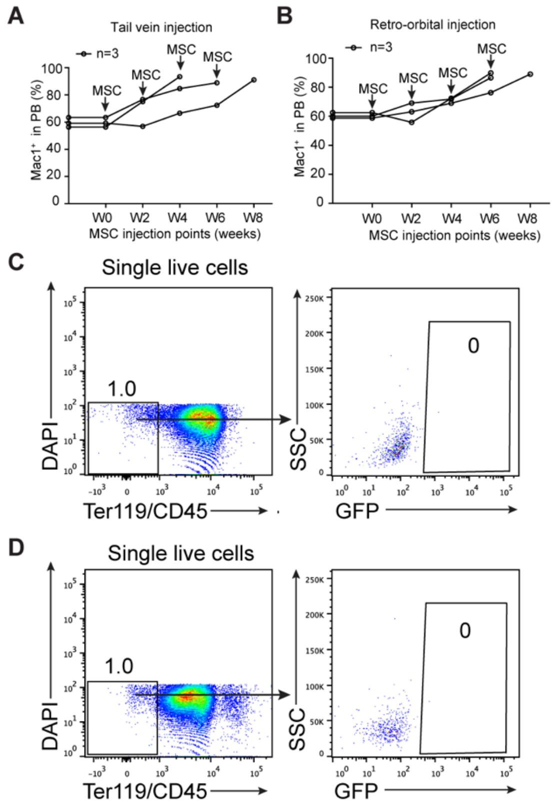


**Figure S6: No therapeutic effect of systemic MSC delivery by tail vein or retro-orbital injection on leukemia-bearing mice**

**(A** and **B)** Kinetics analysis of tumor burden ($CD11b^+$) of MSC-treated leukemia-bearing mice by tail vein injection (A) and retro-orbital injection (B). The time window of MSC treatment is from 0 weeks (W0) to 8 weeks (W8). Flow cytometry analysis of $CD11b^+$ cells in PB was performed every two weeks. A sequential doses of MSCs (2.5 × 10^7 MSCs/kg per dose in 100 µL DPBS) were delivered into each mouse using a 29-gauge needle. N = 3 mice for each group. **(C** and **D)** Flow cytometry analysis indicated that tail vein injection (C) and retro-orbital injection (D) of donor MSCs ($GFP^+$) failed to home to the bone marrow of leukemia-bearing mice. A dose of MSCs (2.5 × 10^7 MSCs/kg in 100 µL DPBS) were delivered into each leukemia-bearing mice using 29-gauge needle by tail vein/retro-orbital injection. One day later, the treated leukemia-bearing mice were sacrificed and their bone marrow cells were analyzed.

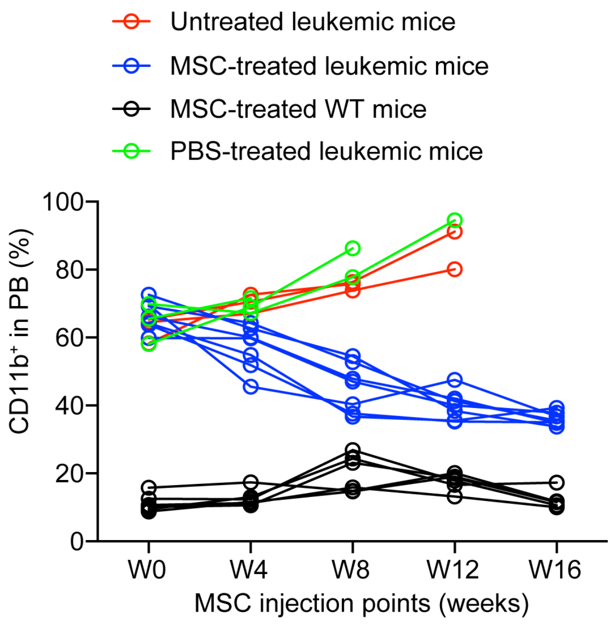


**Figure S7: Intra-BM transfusion of donor MSCs reduces peripheral blood tumor burden of leukemia-bearing mice**

Kinetic analysis of tumor burden ($CD11b^+$) of MSC-treated leukemia-bearing mice. The time window of MSC treatment is from week-0 (W0, treatment starting time) to week-16 (W16). Flow cytometry analysis of tumor burden ($CD11b^+$) in PB was performed monthly. These leukemia-bearing mice were accumulated from multiple independent experiments. The individual leukemia-bearing mice were selected to perform treatment once they reached the leukemia burden standard ($CD11b^+$% > 60% in peripheral blood), but at a different time from multiple experiments. Untreated leukemia-bearing mice were used as disease control (red line), PBS-treated leukemia-bearing mice were used as injection control (green line), and MSC-treated WT mice were used as treatment control (black line). Untreated leukemia-bearing mice: n = 3; PBS-treated leukemia-bearing mice: n = 3; MSC-treated leukemia-bearing mice (blue line): n = 7; MSC-treated WT mice: n = 6.

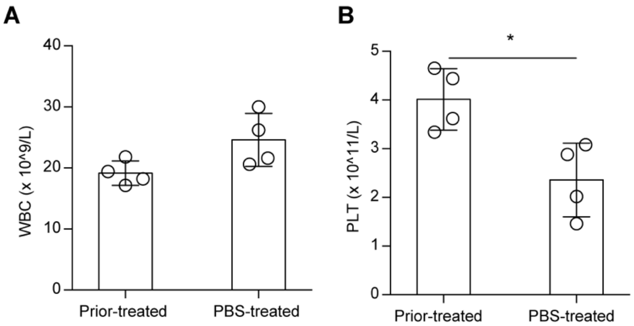


**Figure S8: PBS treatment fails to improve hematopoiesis in leukemic mice**

Statistical analysis of platelets counts in the PB of untreated leukemia-bearing mice and PBS-treated leukemia-bearing mice four weeks after PBS-treatment. **(A)** WBC: white blood cells; **(B)** PLT: platelets. Data are analyzed by unpaired student's t-test (two-tailed). *p < 0.05. Data are represented as mean ± SD (Untreated, n = 4 mice, PBS-treated, n=4 mice).

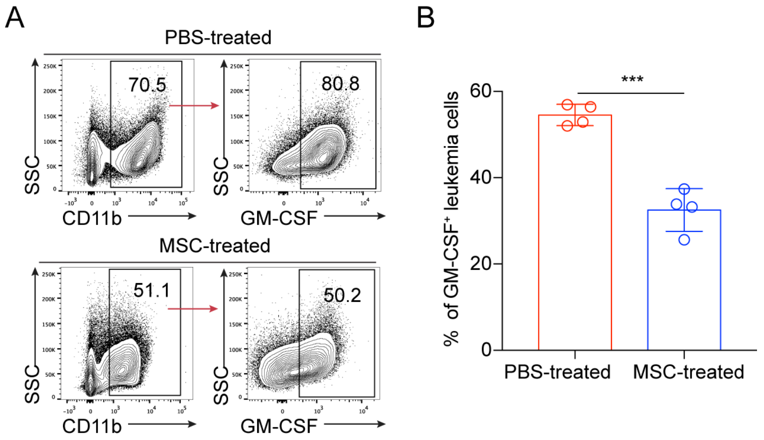


**Figure S9: The expression of GM-CSF proteins in leukemic cells after PBS/MSC treatment**

(A) Representative plots of intracellular flow cytometry staining of GM-CSF proteins in leukemic cells from PBS/MSC-treated leukemia-bearing mice. (B) Statistical analysis of the expression of GM-CSF proteins in leukemic cells. Data are analyzed by unpaired student's t-test (two-tailed). ***p < 0.001. Data are represented as mean ± SD (n = 4 mice for each group).

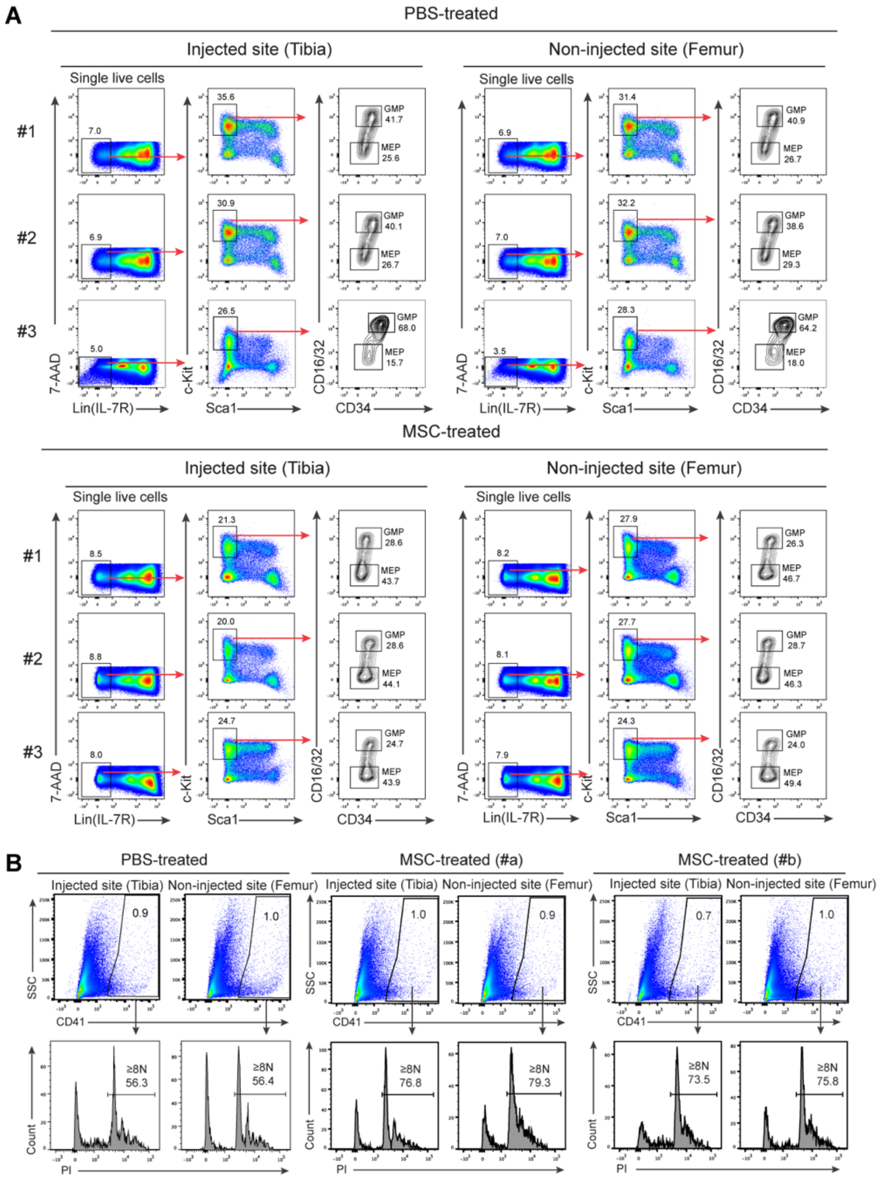


**Figure S10: Systemically re-balanced myeloid lineage progenitor cells and systemically activated megakaryocytes in MSC-treated leukemia-bearing mice**

**(A)** Ratios of myeloid progenitor subpopulations in MSC- and PBS-treated leukemia-bearing mice. Total bone marrow nucleated cells were from the MSC-injected site (tibia) and non-injected site (femur) of each leukemia-bearing mouse, or PBS-injected site (tibia) and non-injected site (femur) of each leukemia-bearing mouse four weeks post MSC/PBS treatment. GMP (granulocyte/macrophage progenitors): $Lin^{-}IL\text{-}7R^{-}Sca1^{-}c\text{-}Kit^{+}CD34^{+}CD16/32^{high}$; MEP (megakaryocyte/erythroid progenitors): Lin$^-$IL-7R$^-$Sca1$^-$c-Kit$^+$CD34$^-$CD16/32$^-$. Three representative plots of PBS-treated leukemia-bearing mice and MSC-treated leukemia-bearing mice four weeks post MSC treatment are shown. **(B)** Activation analysis of megakaryocytes in MSC- and PBS-treated leukemia-bearing mice. Plots from one PBS-treated leukemia-bearing mouse and two MSC-treated leukemia-bearing mice four weeks post MSC treatment are shown. Percentages of mature megakaryocytes with 8N and greater ploidy (≥8N) are shown.

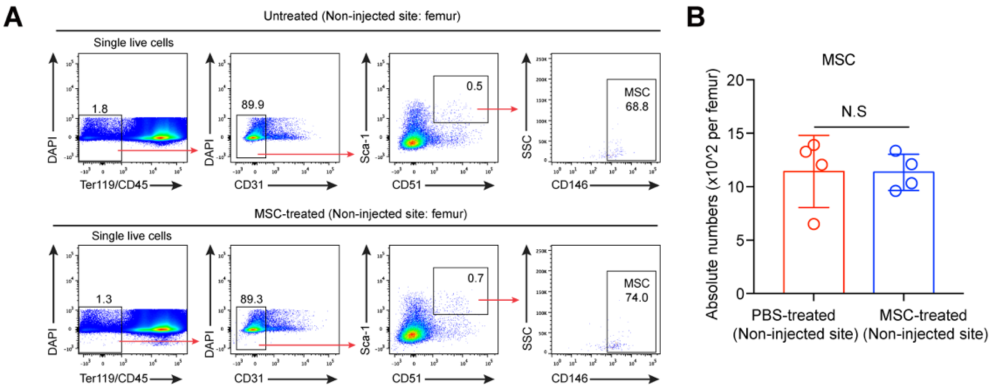


**Figure S11: No improvement in the quantity of host MSCs at the non-injected sites of MSC-treated leukemic mice**

**(A)** Flow cytometry analysis of MSCs at the non-injected site from leukemia-bearing mice eight weeks post MSC treatment. Femurs (non-injected site) of MSC-treated leukemia-bearing mice eight weeks after the first dose MSC treatment were analyzed. Plots of one representative mouse from each group are shown. MSCs are defined as Ter119$^-$CD45$^-$CD31$^-$Sca1$^+$CD51$^+$CD146$^+$. The nucleated cell mixtures of BM and compact bones were prepared for flow cytometry analysis of MSCs. **(B)** Statistical analysis of the absolute numbers of MSCs in the femurs (non-injected site) from untreated and MSC-treated leukemia-bearing mice. Data are analyzed by unpaired student's t-test (two-tailed). N.S indicates not-significant. Data are represented as mean ± SD (n = 4 mice for each group).

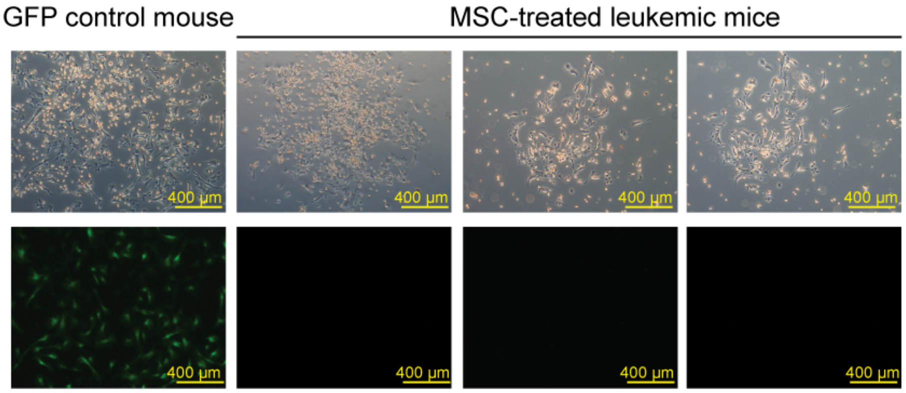


**Figure S12: Recovered MSCs are host origin in MSC-treated leukemia-bearing mice**

Colony-forming units (CFU-F) analysis of functional MSCs from BMNC of control mice (GFP mice) and MSC-treated leukemia-bearing mice. BMNC equivalent to 100 MSCs from control (GFP mice) or MSC-treated leukemia-bearing mice were used as cell input for individual wells (six-well plate). Half-medium change was performed at day 7. Representative fluorescent images of 14-day CFU-F colonies are shown. Bars = 400 µm.

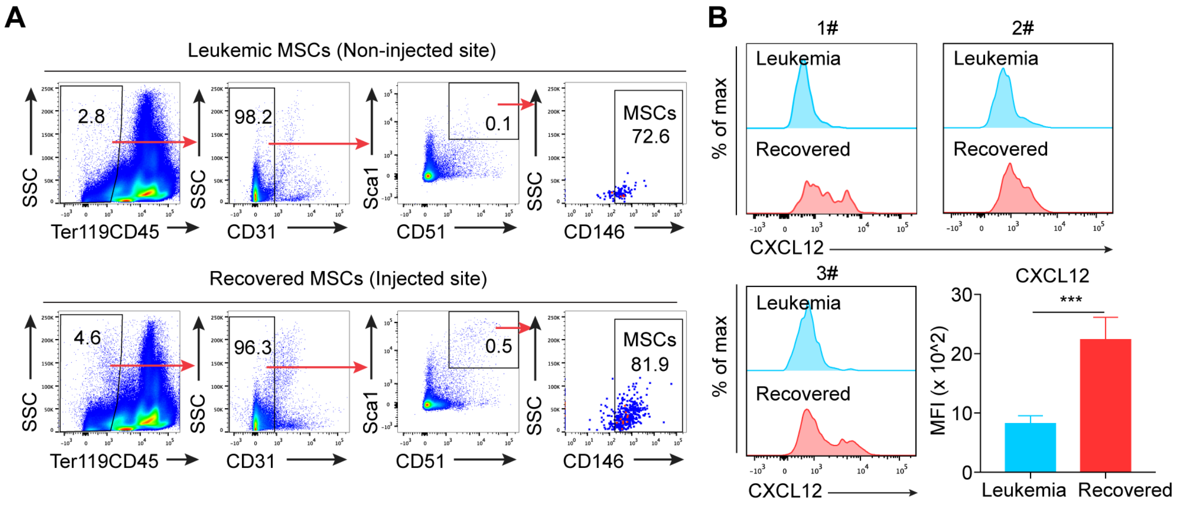


**Figure S13: Restoration of CXCL12 proteins in recovered MSCs**

**(A)** Representative gating strategy of control and leukemic MSCs for intracellular flow cytometry staining analysis of CXCL12 proteins. Plots from one representative tibia (MSC-injected site) and one representative femur (Non-injected site) of MSC-treated leukemia-bearing mouse eight weeks after the first dose MSC treatment are shown. The primary nucleated cell mixtures of BM and compact bones were prepared for intracellular flow cytometry staining. **(B)** Representative intracellular staining plots of CXCL12 proteins in recovered MSCs and leukemic MSCs analyzed by flow cytometry. Statistical analysis of the mean fluorescence intensities (MFI) of CXCL12 in leukemic and recovered MSCs. Data are analyzed by unpaired student's t-test (two-tailed). ***p < 0.001. Data are represented as mean ± SD (n = 6 mice for each group).

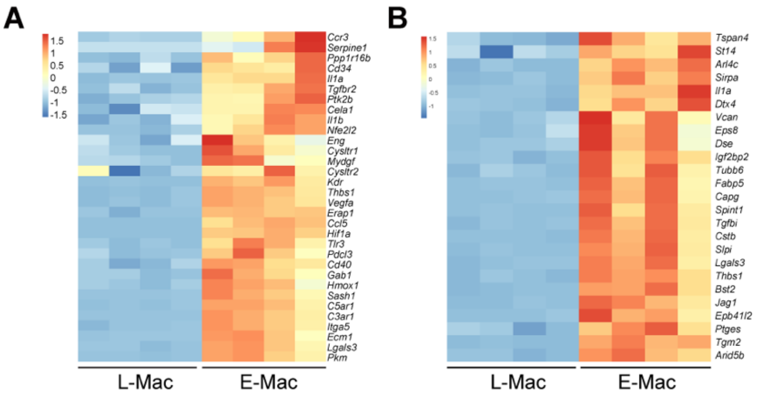


**Figure S14: Up-regulation of leading edge genes related to positive regulation of angiogenesis and cell migration pathways in MSC-reprogrammed macrophages from leukemia-bearing mice**

GSEA of the expression of the leading edge gene subsets. Positive regulation of angiogenesis- **(A)** and cell migration-related **(B)** genes up-regulated in MSC-reprogrammed macrophages from leukemia-bearing mice are shown (a difference in expression of over 2-fold; adjusted p value, < 0.05 (DESeq2 R package). Mac: n = 4 cell sample replicates (one per column); E-Mac: n = 4 cell sample replicates (one per column).

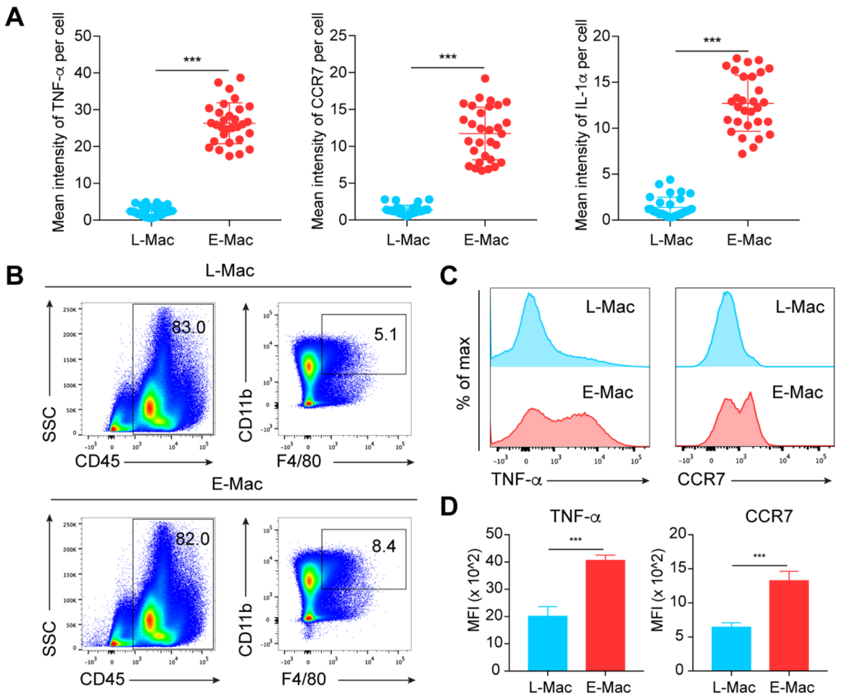


**Figure S15: Elevation of TNF-α, CCR7, and IL-1α in MSC-reprogrammed macrophages at protein level**

**(A)** Statitical analysis of mean intensities of TNF-α, CCR7, and IL-1α fluorescence in L-Mac and E-Mac. Each dot represents a single cell. Data are analyzed by unpaired student's t-test (two-tailed). ***$p < 0.001$. Data are represented as mean ± SD. L-Mac, n=30; E-Mac, n=30. **(B)** Representative gating strategy of L-Mac (Non-injected sites) and E-Mac (MSC-injected sites) in intracellular flow cytometry staining of TNF-α and CCR7 proteins. L-Mac (Non-injected site) and E-Mac (MSC-injected site) were from MSC-treated leukemic mice 12 h post-treatment *in vivo*. **(C)** Representative intracellular staining plots of TNF-α and CCR7 proteins in L-Mac and E-Mac analyzed by flow cytometry. **(D)** Statistical analysis of the mean fluorescence intensities (MFI) of TNF-α, and CCR7 in L-Mac and E-Mac. Data are analyzed by unpaired student's t-test (two-tailed). ***$p < 0.001$. Data are represented as mean ± SD (n = 6 mice for each group).

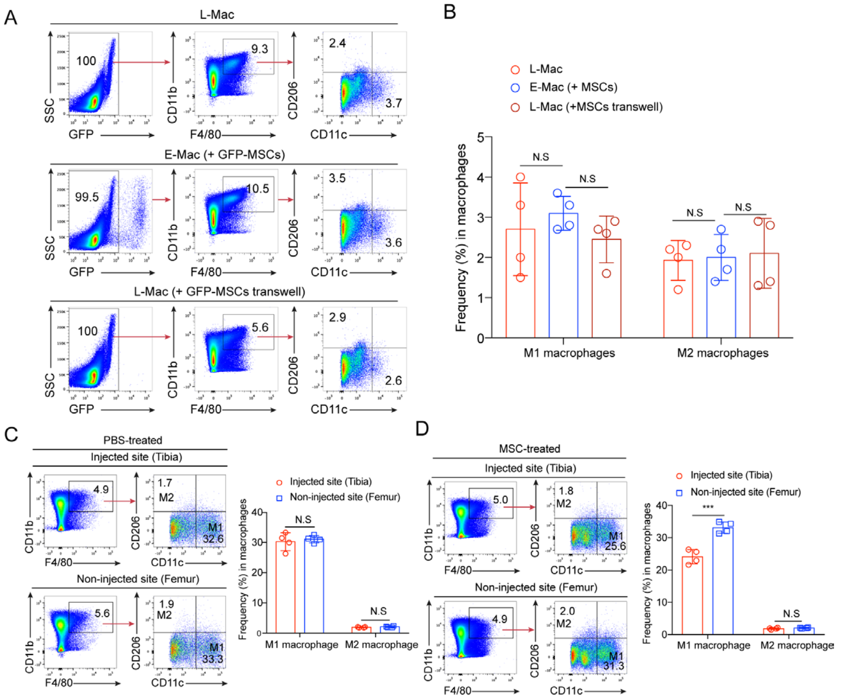


**Figure S16: The MSC-reprogrammed macrophages are not in a manner of phenotypic transformation from M1 into M2**

**(A)** Representative plots of phenotypic M1 and M2 macrophage subpopulations in leukemic macrophages co-cultured with MSCs *in vitro*. M1 macrophages are defined as: CD45$^{+}$CD11b$^{+}$F4/80$^{+}$CD11c$^{+}$CD206$^{-}$, M2 macrophages are defined as: CD45$^{+}$CD11b$^{+}$F4/80$^{+}$CD11c$^{-}$CD206$^{+}$. L-Mac indicates leukemic macrophages. E-Mac (+MSCs) indicates MSC-reprogrammed leukemic macrophages, which were co-cultured with MSCs *in vitro* for 12 h. L-Mac (+MSCs transwell) indicates leukemic macrophages, which were co-cultured in transwell with MSCs *in vitro* for 12 h. **(B)** Statistical analysis of the percentages of M1 and M2 in leukemic macrophages co-cultured with MSCs *in vitro*. Data are represented as mean ± SD (n = 4 mice for each group). **(C)** Representative plots of phenotypic M1 and M2 macrophage subpopulations from PBS-/MSC-treated leukemia-bearing mice. M1 macrophages are defined as: CD45$^{+}$CD11b$^{+}$F4/80$^{+}$CD11c$^{+}$CD206$^{-}$, M2 macrophages are defined as:

$CD45^+CD11b^+F4/80^+CD11c^-CD206^+$. The bone marrow nucleated cells of PBS-/MSC-injected sites (tibias) and non-injected sites (femurs) from leukemia-bearing mouse 12 h post-treatment were prepared for flow cytometry analysis of macrophage subtypes. **(D)** Statistical analysis of the percentages of M1 and M2 macrophages. Data are analyzed by one-way ANOVA test. ***$p < 0.001$, N.S indicates not-significant. Data are represented as mean ± SD (n = 4 mice for each group).

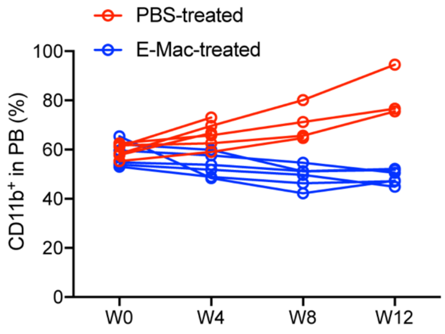


**Figure S17: Kinetic analysis of tumor burden ($CD11b^+$) of PBS/E-Mac treated leukemia-bearing mice**

PBS-treated leukemia-bearing mice were used as injected control. *$p < 0.05$. Data are analyzed by one-way ANOVA test. Data are represented as mean ± SD. PBS-treated: n = 6 mice; E-Mac-treated: n = 6 mice.

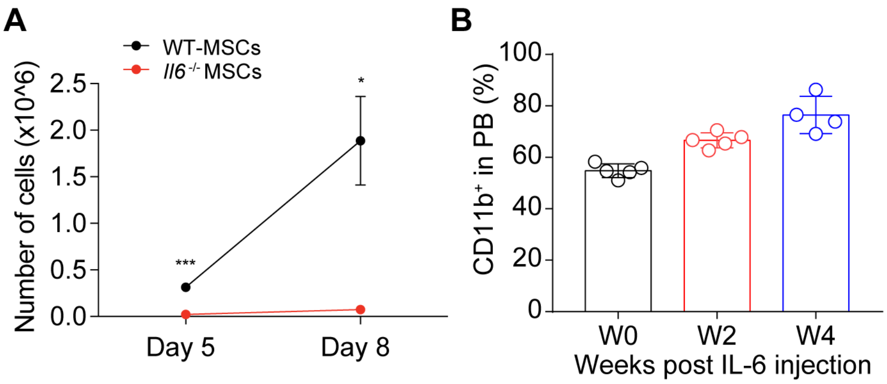


**Figure S18: Systemic transfusion of IL-6 proteins fails to suppress leukemia**

**(A)** The primary MSCs were isolated from the compact bones and bone marrow nucleated cells (BMNCs) of 3-4 weeks old healthy WT mice and *Il6*$^{-/-}$ mice. The primary bone marrow MSCs equivalent to each mouse were used as input for proliferation in α-MEM medium containing 10% FBS. The absolute numbers of cells were counted before passaging on day 5 and day 8. Data are analyzed by unpaired student's t-test (two-tailed). *P < 0.05, ***P < 0.001. Data are represented as mean ± SD (n = 3 independent experiments). **(B)** Kinetics analysis of tumor burden (CD11b$^{+}$) of IL-6-treated leukemia-bearing mice. Flow cytometry analysis of CD11b$^{+}$ cells in PB was performed every two weeks. A dose of 40 μg/kg IL-6 proteins were injected into each leukemia-bearing mouse by intraperitoneal injection, and the injection interval is 24 hours. Data are represented as mean ± SD (n = 5 mice).

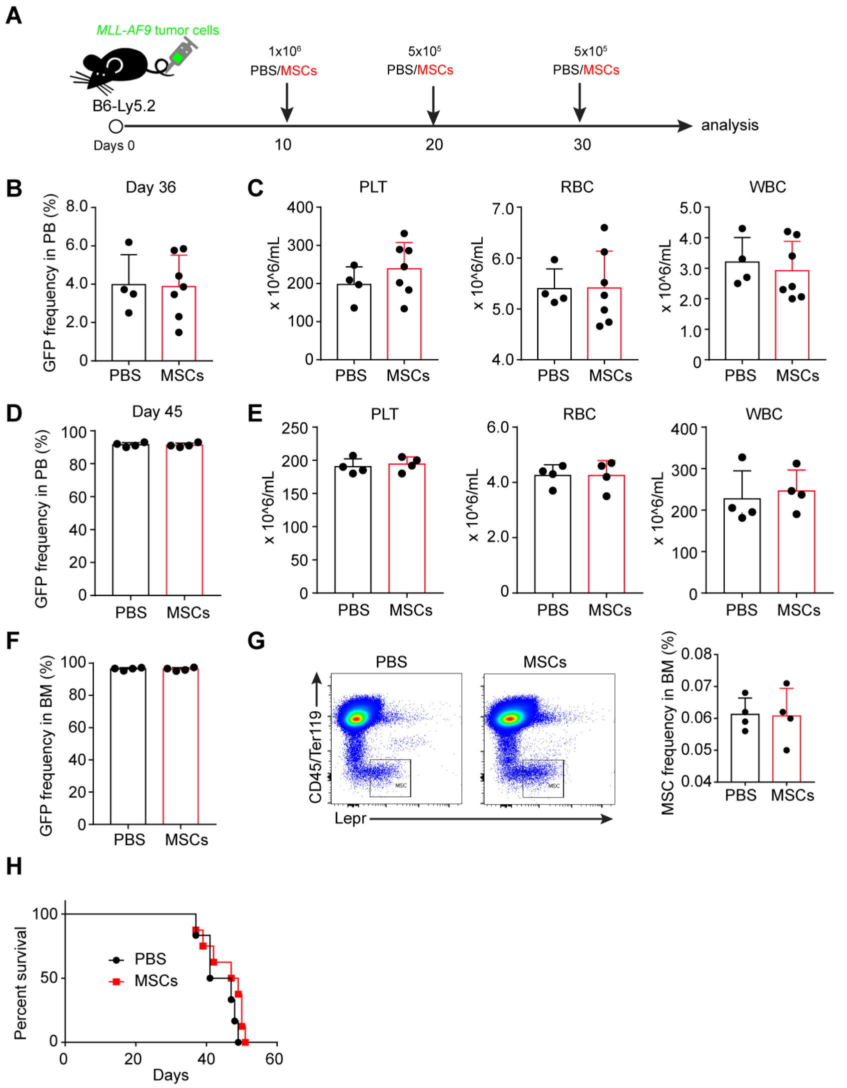


**Figure S19: MSC treatment fails to suppress AML initiated by *MLL-AF9* translocation**

**(A)** Treatment protocol. AML cells (GFP$^{+}$) were i.v. injected at day 0, and then the mice were intra-tibia injected with 10 μL PBS or MSCs every 10 days from day 10. **(B)** Frequency of AML cells in PB at day 36. **(C)** PLT, RBC and WBC counts of AML-bearing mice at day 36. **(D)** Frequency of AML cells in PB at day 45. **(E)** PLT, RBC and WBC counts of AML-bearing mice at day 45. **(F)** Frequency of AML cells in BM at day 45. **(G)** Flow cytometry analysis of frequencies (right panel) of MSCs in the BM of leukemia-bearing mice at day 45. **(H)** Survival curve of AML-bearing mice treated with PBS or MSC. Error bars show SD, n=4-6.

### References

1. Cheng H, Hao S, Liu Y, et al. Leukemic marrow infiltration reveals a novel role for Egr3 as a potent inhibitor of normal hematopoietic stem cell proliferation. *Blood*. 2015;126(11):1302-1313.

2. Zhu H, Guo ZK, Jiang XX, et al. A protocol for isolation and culture of mesenchymal stem cells from mouse compact bone. *Nat Protoc*. 2010;5(3):550-560.
